# Converging cytokine and metabolite networks shape asymmetric T cell fate at the term human maternal-fetal interface

**DOI:** 10.1101/2024.06.10.598377

**Authors:** Nicholas J Maurice, Jami R Erickson, Caitlin S DeJong, Florian Mair, Alexis K Taber, Marie Frutoso, Laura V Islas, Anna-Lena BG Vigil, Richard L Lawler, M Juliana McElrath, Evan W Newell, Lucas B Sullivan, Raj Shree, Stephen A McCartney

## Abstract

Placentation presents immune conflict between mother and fetus, yet in normal pregnancy maternal immunity against infection is maintained without expense to fetal tolerance. This is believed to result from adaptations at the maternal-fetal interface (MFI) which affect T cell programming, but the identities (i.e., memory subsets and antigenic specificities) of T cells and the signals that mediate T cell fates and functions at the MFI remain poorly understood. We found intact recruitment programs as well as pro-inflammatory cytokine networks that can act on maternal T cells in an antigen-independent manner. These inflammatory signals elicit T cell expression of co-stimulatory receptors necessary for tissue retention, which can be engaged by local macrophages. Although pro-inflammatory molecules elicit T cell effector functions, we show that additional cytokine (TGF-β1) and metabolite (kynurenine) networks may converge to tune T cell function to those of sentinels. Together, we demonstrate an additional facet of fetal tolerance, wherein T cells are broadly recruited and restrained in an antigen-independent, cytokine/metabolite-dependent manner. These mechanisms provide insight into antigen-nonspecific T cell regulation, especially in tissue microenvironments where they are enriched.

## Introduction

Despite having evolved under the pressures of adaptive immunity, placentation is immunologically paradoxical in which maternal immunity tolerates a fetal allograft while maintaining competency towards other pathogens(1). By virtue of thymic selection, maternal T cells are selected for reactivity to non-self peptide antigens (Ag) sourced from intracellular protein degradation(2); but half of the fetal genome encodes paternal alleles that could be seen as non-self. Previous studies have illustrated that the placenta, the component of the fetal graft most exposed to maternal immunity, has multiple mechanisms to avoid detection by the maternal immune system: placental trophoblasts downregulate the expression of major histocompatibility complex class I (MHC-I), the molecule required for surface presentation of Ags, to limit maternal T cell recognition and killing(3). Further studies have found additional evasive tactics used to restrict maternal T cells at the site of placentation (i.e., the maternal-fetal interface: MFI) including decreased perforin and granzyme production and increased expression of co-inhibitory molecules compared to T cells from the blood(4–6). Mouse models found that the decidua (the endometrial tissue which is remodeled by invading placental trophoblast), forbids maternal T cell entry by silencing the expression of chemokines encoded by *Cxcl9*, *Cxcl10*, and *Ccl5*, even during artificially high levels of systemic inflammation(7). Together, these data suggest that maternal immunocompetency and fetal tolerance is achieved by mechanisms that make the MFI as immunologically invisible as possible.

Despite these mechanisms, there is mounting evidence that the MFI, and pregnancy itself, are not immunologically silent. Early in pregnancy, natural killer (NK) cells at the decidua contribute to effective placentation(8–12); and NK cells, as well as T cells, can be found at the first-trimester MFI using single-cell RNA sequencing (scRNAseq)(13). Pregnancy affects T cells, elevating regulatory T cell (Treg) frequencies(14–16), increasing the frequency of T cells at the MFI as pregnancy progresses, including clones with specificity to fetal Ag systemically(17, 18). As pregnancy progresses, so does a signature of IL-2–STAT5 signaling, which ceases postpartum(19). At the end of gestation, signatures of immune cell activation can be detected in human choriodecidual tissues(20), suggesting immunity’s role in both pregnancy and labor. Furthermore, inducing inflammation at the MFI with T cell receptor (TCR) or Toll-like receptor (TLR)-4 agonists, can lead to preterm labor in mice(21). Given the immunologic nature of these phenomena, this suggests that maintaining homeostasis during maternal-fetal conflict is not addressed by evasion alone, and likely involves selective tuning of immune cell functions, but the details of how these pro- and anti-inflammatory signals are balanced remains poorly understood.

Cytokines have the potential to activate and reshape CD8+ T cell fates even in the absence of cognate Ag. This phenomenon, called bystander activation, partially mirrors that of TCR-mediated activation: bystander-activated T cells can proliferate, express cytokines, and directly kill targets in an antigen-independent, inflammation-dependent manner(22, 23). Indeed, fetus-specific and bystander (i.e., virus-specific) CD8+ T cell populations at the MFI appear similarly activated, suggesting T cell activation at the MFI that is agnostic to TCR specificity(18). Though significant alterations to pro- and anti-inflammatory cytokines occur during pregnancy, these are understood from systemic (i.e., peripheral serum) measurements(24, 25). Thus, the specific activating and inhibitory signals involved in locally (i.e., at the MFI) regulating CD8+ T cell fate during pregnancy is incompletely defined.

Using samples from a cohort of patients undergoing uncomplicated term cesarean deliveries, we used high-parameter flow cytometric and untargeted transcriptomic single-cell analyses to uncover maternal T cell adaptations unique to the MFI. We found a subset of CD8+ T cells are tissue resident at the MFI and highly activated. Despite activation signatures congruent with TCR activation, we find this phenomenon is likely independent of TCR specificity or cognate Ag. Using a combination of single-cell transcriptomics and proteomics, we instead discover pro-inflammatory chemokine and cytokine networks which may recruit and activate maternal CD8+ T cells at the MFI. Though these signals are sufficient to elicit CD8+ T cell cytotoxicity in vitro, we find this is likely curbed by elevated levels of regulatory molecules in situ. Specifically, we find elevated transforming growth factor-β1 (TGF-β1) and the tryptophan metabolite, kynurenine, at the placenta. During in vitro cytokine stimulation, these factors cooperate to limit inflammation-mediated cytotoxicity without restricting other programs resulting from inflammation-dependent activation. Together, our data suggests that mechanisms at the MFI recruit, retain, and restrain maternal T cells at the maternal side of the MFI. Importantly, these factors at the placenta do not attenuate TCR-driven CD8+ T cell effector programs in vitro, suggesting that Ag-specific immunity is not compromised by elevated TGF-β1 or kynurenine. We discuss the relevance of inflammation-dependent CD8+ T cell programs for both fetal tolerance and maternal competence against pathogens.

## Results

### A tissue-resident phenotype is biased towards the maternal side of the MFI

The maternal-fetal interface (MFI) is the primary site of immune conflict, so we initially sought to comprehensively test changes in cellular immunity here versus the periphery of mother and fetus. Since our initial aim was to develop a dataset concerning immunity at the MFI during homeostasis at term, we selectively recruited participants without pregnancy and/or immunologic complications undergoing routine, term (37-41 weeks gestation), unlabored cesarean section. We chose to exclude cases undergoing vaginal delivery or cesarean after labor to minimize immune modifications that have been associated with labor and parturition(26). Since the placenta is a circulatory interface, we isolated leukocytes from both maternal blood (MB) and fetal cord blood (CB), to determine if our observations resulted from peripheral cell contamination or tissue-dependent phenomena (**Figure 1A**). The fetal-derived placenta is heterogenously exposed to maternal immunity; therefore, we sampled multiple biopsies from the maternal-facing side of the placenta (including adherent decidual tissue) and the fetal-facing side of the placenta (mPLAC and fPLAC, respectively) (**Figure 1A**). We interrogated these tissues using 28-color flow cytometry(27), including cryopreserved peripheral blood mononuclear cells (PBMC) from a single non-pregnant Seattle Area Cohort (SAC) donor to determine batch-to-batch variation (**Figure 1A**).

**Figure 1.**
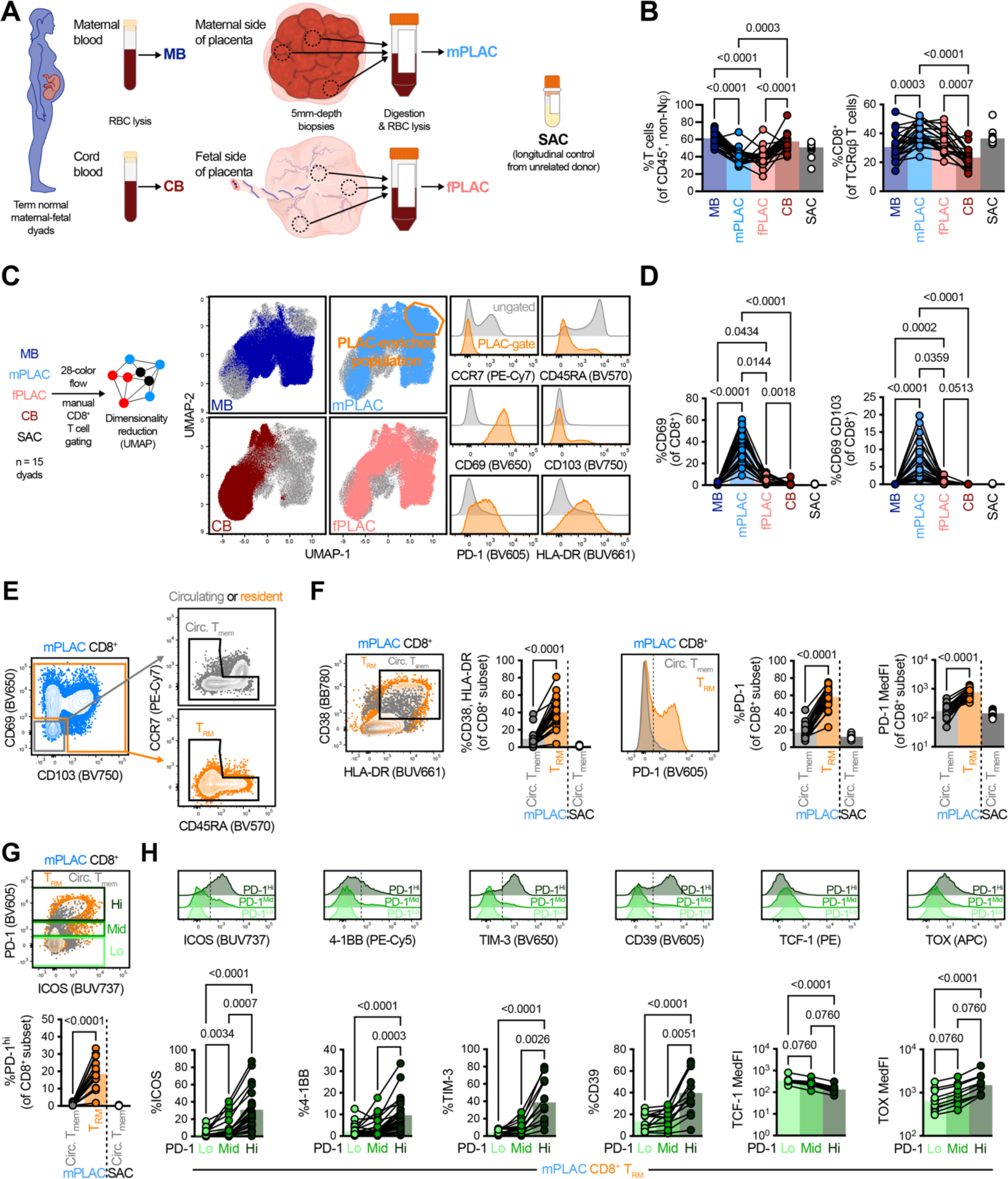
A T_RM_ population with an activated phenotype is asymmetrically distributed at the human MFI **A** Overview of tissue sampling for cytometric analyses of leukocytes in maternal blood (MB), cord blood (CB), the maternal or fetal sides of the MFI (mfPLAC and fPLAC, respectively), and a longitudinal PBMC donor from the Seattle Area Cohort (SAC). **B** Frequencies of bulk and CD8^+^ T cells across tissues. **C** 28-color T cell phenotyping flow data for CD8^+^ T cells from 15 dyads visualized by UMAP and subsequent identification of a PLAC-enriched subpopulation. **D** Frequency of T_RM_-phenotype (i.e. CD69^+^ and CD69^+^CD103^+^) CD8^+^ T cells across tissues. **E** Representative gating of CD8^+^ T_RM_ and circulating T_mem_ from mfPLAC samples. **F** Comparisons of activation markers in CD8^+^ T_RM_ and circulating T_mem_. **G** Frequency of PD-1^hi^ events within CD8^+^ subsets and representative gating. **H** Activation phenotypes across mfPLAC CD8^+^ T_RM_ stratified by PD-1 intensity. **B** depicts 22-25 dyads, C depicts 10,000 CD8^+^ T cells per tissue, per dyad from 15 dyads. **D** depicts 20-21 donors. **G** depicts 21 donors. **H** depicts 10–25 donors. All symbols represent a unique population which are connected across donor identity with bars depicting mean. All statistical significances where *p* < 0.1 are indicated as calculated by **B**, **D**, **H** Friedman tests and Dunn’s multiple comparison tests or **F**, **G** Wilcoxon tests.

Though we found a decreased frequency of CD3+ T cells among leukocytes in placental tissues, CD8+ T cells constituted a greater proportion of the T cell pools at these sites than in circulation (**Figure 1B**). Given the difference in CD8+ T cell frequency, we surveyed for other tissue-based changes in phenotypes. Owing to the high complexity of 28-color flow data, we first utilized dimensionality reduction using uniform manifold approximation projections (UMAP)(28) on pre-gated CD8+ T cell events (**Figure 1C**). When we color-coded CD8+ T cell events based on tissue source, we noted a population dominated by cells isolated from mPLAC biopsies (**Figure 1C** PLAC-enriched gate). This population, compared to all other events, was characterized by T cells with an effector memory (TEM) phenotype (CCR7lo, CD45RAlo) and upregulation of markers associated with activation, exhaustion, and/or tissue residence (CD69, CD103, PD-1, and HLA-DR) (**Figure 1C, Supplemental figure 1A, B**).

Previous studies have described an enrichment of activated (i.e., CD69-expressing) and/or TEM at the MFI(29, 30), yet these studies predated the discovery of tissue resident memory (TRM) cells and biomarkers which define them(31). TRM share a TEM phenotype (CCR7lo CD45RAlo), yet near-constitutively express the activation marker CD69 and reside in tissues long-term(32, 33). Therefore, we asked if PLAC-enriched CD8+ T cells were of a TRM phenotype. We surveyed the expression of CD69 and CD103, which can be singly or doubly expressed by TRM (31, 32, 34). Despite donor-to-donor variation, we found significant expression of CD69 in mPLAC CD8+ T cells, and to a lesser degree, CD69 CD103 co-expression (**Figure 1D**). Further, nearly all CD69+ events from mPLAC CD8+ T cells maintained a phenotype similar to those of TEM (CCR7lo, CD45RAlo, **Supplemental figure 1C**), in line with using CD69 as a biomarker of tissue residence rather than activation alone.

### CD8+ TRM populations are maternally derived and highly activated

To ensure appropriate use of CD69 as a biomarker of tissue residence, we leveraged single-cell RNA sequencing (scRNAseq) on CD8+ T cells that we sorted fresh from tissues using fluorescence-activated cell sorting (FACS). Like our targeted cytometry-based proteomics data (**Figure 1C**), we could identify a placenta-enriched subpopulation of CD8+ T cells using untargeted transcriptomics (**Supplemental figure 2A**). We identified this population with graph-based clustering (cluster 5) and found it had transcriptional profiles concurrent with mouse and human CD8+ TRM from other organs (low *CCR7*, *SELL*, *KLF2*, *KLF3*, *TCF7*, *S1PR4*; high *CD69*, *RGS1*) (Supplemental figure 2A, B)(34–37). Since mother (XX) and fetus (XY) were sex mismatched, we were able to identify this TRM population as maternal-derived, as it expressed higher levels of the X-inactivating transcript, *XIST*, and negligible levels of the Y-associated transcript, *RPS4Y1* (**Supplemental figure 2B**). To further prove this phenotype was one of bona fide tissue residence, rather than transient activation of circulating CD8+ T cells in the placenta, we isolated maternal intervillous blood (IVB) for flow analysis (**Supplemental figure 2C**). IVB CD8+ T cell frequency and phenotype mirrored that of maternal circulation, indicating a non-circulating TRM population. Indeed, we found CD8+ TRM to be associated with the decidua basalis, a thin layer of remodeled endometrial tissue that remains adhered to the placenta due to interdigitation with invading placental cytotrophoblasts (**Supplemental figure 2D**).

The decidua basalis is the tissue site at which maternal immunity is most exposed to the fetal allograft. Having found CD8+ TRM at the MFI, we asked if these cells have additional modifications in comparison to memory T cells (Tmem) that circulate within this tissue (i.e., CD69- CD103- cells). We chose to omit naïve (i.e., CCR7+ CD45RA+) CD8+ T cells from this comparison (**Figure 1E**) since naïve T cells do not form tissue residence(32, 38) and are unable to respond to the same breadth of stimuli as Tmem (whether resident- or circulating-memory)(22). In comparison to their circulating counterparts, CD8+ TRM were more frequently of an activated phenotype, expressing higher percentages of HLA-DR and CD38 or PD-1 (**Figure 1F**). Surprisingly, mPLAC CD8+ TRM expressed high levels of PD-1 (**Figure 1G**), which is often only seen after TCR-mediated activation or dysfunction(39, 40). We therefore sought to determine whether PD-1 expression levels in mPLAC CD8+ TRM correlated with other phenotypes reflecting TCR stimulation, including ICOS(41), 4-1BB(42), TIM-3(43, 44), and CD39(45). A considerable fraction of PD-1hi CD8+ TRM co-expressed these proteins (**Figure 1H, Supplemental figure 1D**). Further, we found altered levels of transcription factors TCF-1 and TOX (**Figure 1H**), paralleling patterns found in T cells rendered dysfunctional by chronic TCR stimulation(46–49). Given these profiles, we asked whether we could determine the TCR specificity of these cells.

### Virus-specific bystander TRM at the MFI are found at frequencies reflecting those in circulation

Since we observed a subset of maternal-derived CD8+ TRM expressing markers indicative of TCR-mediated activation and/or exhaustion, we sought to test their TCR specificities. We identified four mothers for whom we had cryopreserved samples (paired MB and mPLAC) for combinatorial tetramer screening using cytometry by time of flight (CyTOF)(50) (**Figure 2A**). These donors specifically encoded HLA alleles compatible for screens via qPCR (HLA-A*01, - A*02, -A*03, -A*011, and/or B*07); our tetramers included Y-associated(17), tumor-associated(45), and viral antigens (Ag) (**Supplemental tables 10, 11**). We specifically gated resident (CD69+ and/or CD103+) and circulating (CD69- CD103-) CD8+ T cells with a memory phenotype (i.e., not CCR7+ CD45RA+) for our analysis (**Figure 2B**). Although we failed to identify CD8+ T cells with specificities for published Y or tumor Ags, we found CD8+ T cells specific for both chronic (Epstein-Barr virus, EBV; cytomegalovirus, CMV; herpes simplex virus, HSV) and acute viruses (influenza A virus, IAV; adenovirus, AdV) (**Figure 2C**). We observed the frequency of CD8+ TRM with defined TCR specificities was proportional to their abundance in the circulating memory pool (in both peripheral or mPLAC) (**Figure 2C**). This notably contrasts with previous studies demonstrating elevated frequencies of virus-specific CD8+ T cells at the MFI(51), which was likely inflated compared to blood due to the inclusion of naïve CD8+ T cells in blood sample analyses, despite their inability to seed tissues.

**Figure 2.**
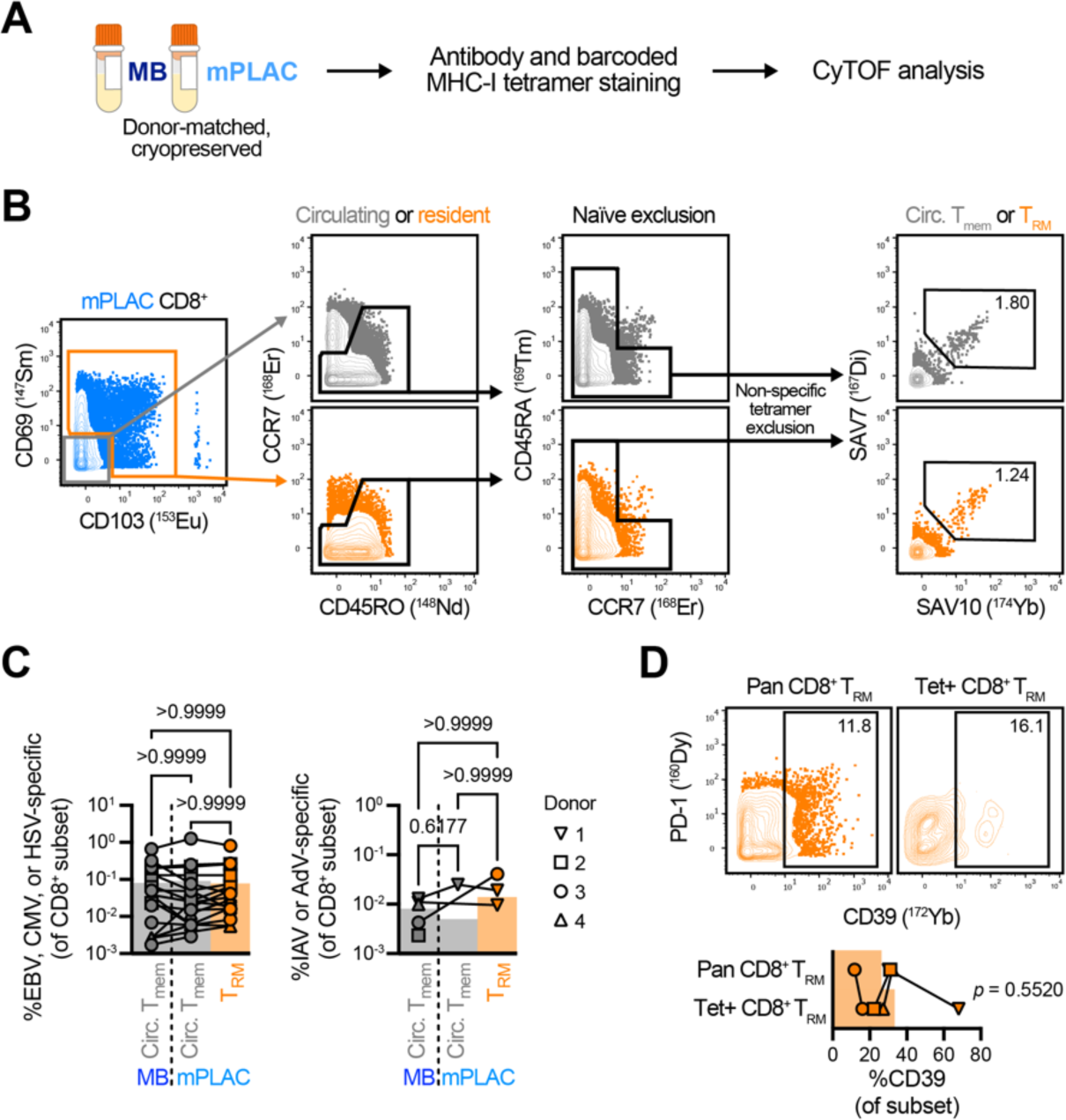
Ag-nonspecific CD8^+^ T_RM_ frequencies at MFI mirror those of circulating CD8^+^ T_mem_ **A** Overview of barcoded tetramer screening of cryopreserved, donor-matched MB and mfPLAC samples. **B** Representative gating of CD8^+^ T_RM_ and circulating T_mem_ populations and downstream tetramer gating. To exclude events that bound MHC-I tetramers in a non-specific manner, we gated events negative for all but two of our tetramers of interest, before identifying events staining positive for our tetramers of interest (here, SAV7 and SAV10 for HLA*11 EBV EBNA4-specific T cells). **C** Frequency of CD8^+^ T cells (circulating T_mem_ from MB, circulating T_mem_ and T_RM_ from mfPLAC) that are specific for chronic/latent viral infections (left; EBV, CMV, VZV, HSV) or acute viral infections (right; IAV, AdV). **D** Frequency of CD39 expression within bulk, polyclonal (pan) and virus specific (tetramer+) CD8^+^ T_RM_. **C** depicts 7, 4, 7, and 3 T cell populations specific for EBV, CMV, or HSV Ag and 1, 1, 1, and 2 T cell populations specific for IAV and AdV Ag from *n* = 4 donors. **D** depicts *n* = 4 donors. Symbols are connected by (**C**) tetramer specificity and donor identity or (**D**) donor identity. Indicated statistical significances were calculated by **C** Friedman tests with Dunn’s multiple comparisons or **D** paired *t* test.

Given the presence of virus-specific “bystander” CD8+ TRM at the MFI, we asked if they fail to upregulate markers of TCR engagement due to their lack of Ag specificity. We chose to interrogate CD39, a biomarker recently found to be elevated on cells receiving in situ TCR stimulation(45). But virus-specific (i.e., tetramer-positive) CD8+ TRM expressed similar levels of CD39 to their bulk counterparts that we could not characterize by tetramer stains (**Figure 2D**). Though we cannot formally exclude alloreactivity of mPLAC CD8+ TRM(52), we believe this to be unlikely since TCR specificities in this population are not dramatically enriched over their frequency in circulating CD8+ Tmem, as one may expect following TCR-mediated expansion. Therefore, we hypothesized that generalized signals (e.g., cytokines/chemokines) may influence TRM fate and phenotypes at the MFI.

### Signaling networks at the placenta predispose for T cell recruitment and retention

We first analyzed untargeted scRNAseq data from donor-matched FACS-enriched CD8+ T cells and non-B cell HLA-DR+ Ag-presenting cells (APCs) with the NicheNet package(53). We specifically used NicheNet to identify ligand-receptor interactions that were intact at the transcriptomic level between APC and CD8+ T cells, querying for interactions that were elevated in mPLAC versus MB populations, as well as those elevated in mPLAC CD8+ TRM versus mPLAC circulating CD8+ Tmem (**Figure 3A, Supplemental figure 3A**). We delineated mPLAC CD8+ TRM from mPLAC circulating CD8+ Tmem as events with a TEM phenotype and high *RGS1* levels. We found that chemokine and cytokine interactions were predicted to be elevated at the mPLAC versus MB (**Supplemental figure 3A**); but that this was not due to anatomic location alone, for these same signaling pathways were predicted to be elevated in mPLAC CD8+ TRM versus mPLAC circulating CD8+ Tmem interacting with the same APCs (**Figure 3B**).

**Figure 3.**
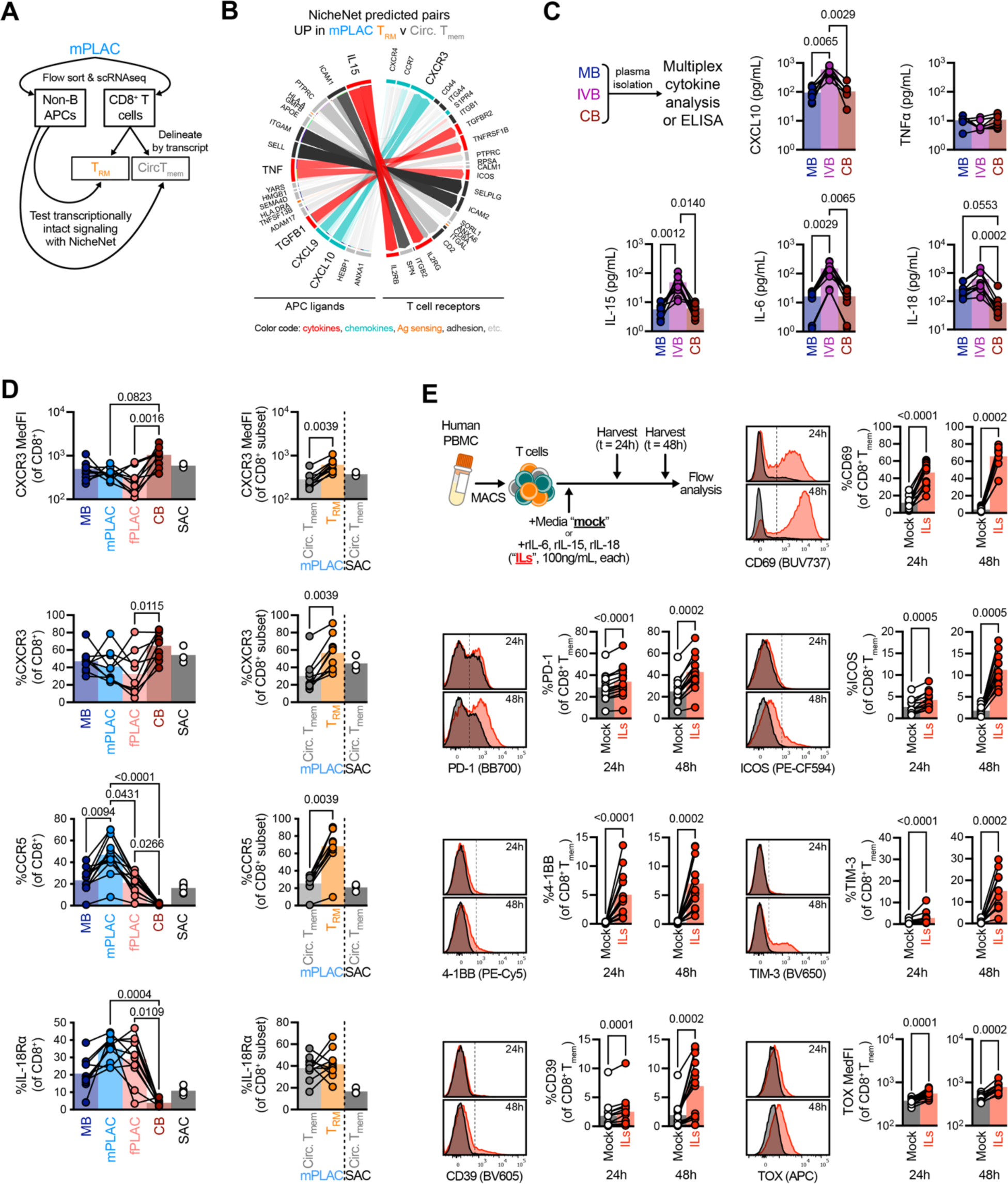
Intact signaling networks at the MFI induce T_RM_-like phenotypes in vitro **A** Experimental approach to identify cellular interactions in mfPLAC. Specifically, we FACS-sorted APCs and CD8^+^ T cells from mfPLAC, conducted single-cell RNAseq, and tested potential ligand-receptor interactions using NicheNet. Non-B cell, HLA-DR^+^ cells were used as the “sender” population and circulating memory and tissue resident CD8^+^ T cell subsets were delineated by transcript and used as the “receiver populations.” **B** Circos plot identifying potential ligand-receptor pairs up in mfPLAC CD8^+^ T_RM_ versus circulating CD8^+^ T_mem_. Interactions implicating cytokines, chemokines, Ag presentation/sensing, and adhesion molecules are respectively colored red, teal, orange, and dark grey. **C** Chemokine and cytokine concentrations determined by multiplex analysis of plasma from MB, IVB, and CB. **D** Expression on chemokine and cytokine receptors in CD8^+^ T cells across tissues (left) and within circulating CD8^+^ T_mem_ and T_RM_ subsets from mfPLAC samples (right). **E** Induction of activation markers with cytokines. Specifically, we MACS-isolated T cells from PBMC samples and stimulated cells with media alone (“mock”) or IL-6, IL-15, and IL-18 in combination (“ILs, ”100ng/mL, each) and measured changes in CD8^+^ T_mem_ phenotypes using flow. **B** depicts scRNAseq data from *n* = 1 dyad with an XY fetus. **C** depicts *n* = 7–9 dyads. **D** depicts *n* = 10–13 dyads. **E** depicts *n* = 12-15 donors across 6 technical replicates. Indicated are statistical significance where *p* < 0.1 by **C**, **D** Friedman tests with Dunn’s multiple comparisons and **C**, **D** Wilcoxon tests.

Since cytokine signaling is in line with our hypothesis of generalized signaling pathways, we next tested if these molecules are indeed expressed at the MFI. We interrogated cytokine and chemokines in MB, CB, and IVB plasma as well as mPLAC and fPLAC tissue lysate supernatants using multiplexed cytokine analysis. We found the chemokine, CXCL10, which can recruit Ag-specific and -nonspecific CD8+ Tmem through CXCR3(54, 55), was elevated in both IVB plasma and tissue lysates (**Figure 3C, Supplemental figure 3B**). Although TNF signaling was intact at the transcriptional level, we were unable to find it elevated at the MFI (**Figure 3C**), illustrating limitations of transcriptional analysis alone(56). Despite this, NicheNet accurately predicted intact IL-15 signaling networks, which like CXCL10, were elevated in IVB plasma and mPLAC lysates (**Figure 3C, Supplemental figure 3B**). IL-15, amongst other cytokines (IL-2, IL-4, IL-7, IL-9, and IL-21) signal through receptor complexes involving the common γ chain (γc), leading to downstream STAT5 signaling. Though a previous study described an immunologic STAT5 signature of pregnancy, hypothesized to result from IL-2 signaling(19), we were unable to find elevated levels of other γc cytokines in IVB plasma or placental tissue lysates (**Supplemental figure 3C–D**). While NicheNet assisted in identifying CXCL10 and IL-15, we also found increased concentrations of IL-6 and IL-18 at the MFI that NicheNet had failed to predict (**Figure 3C, Supplemental figure 3B**). This could result from these signals originating from a different cell type (i.e., non APCs) at the MFI or to limitations in sensitivity with smaller, non-targeted scRNAseq datasets(56). Using flow cytometry, we found that CD8+ T cells express factors underlying their recruitment and cytokine mediated activation (**Figure 3D**) suggesting that these signaling pathways can indeed influence CD8+ Tmem at the MFI.

We next sought to determine how pro-inflammatory cytokines at the MFI could augment T cells in vitro. We specifically cultured T cells with IL-6, IL-15, and IL-18 for up to 48h (**Figure 3E**) and then assessed cell phenotype. We specifically enriched T cells from cryopreserved PBMCs with negative selection magnet-activated cell sorting (MACS) to mitigate indirect activation that could arise from contaminating APCs. While pro-inflammatory cytokines sufficiently activated CD8+ Tmem (vis-à-vis CD69 upregulation), we also saw induction of proteins observed in CD8 TRM from mPLAC biopsies, including PD-1, ICOS, 4-1BB, TIM-3, CD39, and TOX (**Figure 3E**).

Therefore, it is possible that pro-inflammatory networks at the MFI recruit and activate CD8+ Tmem even in the absence of cognate Ag. Though we observed induction of co-stimulatory and-inhibitory receptors on T cells, these receptors must engage ligands to alter T cell fates, which can include tissue retention (57, 58). We hypothesized that co-stimulatory and -inhibitory receptors that are elevated on CD8+ TRM at the MFI could be engaged by ligands on other cells at this site.

### Decidual macrophages express cognate ligands for inflammation-induced receptors and physically engage CD8+ T cells in situ

We tested which APC subset was most likely influencing NicheNet predictions and found that CD14+ macrophages/monocytes were the sole APC subset overrepresented within mPLAC versus MB (**Figure 4A, Supplemental figure 4A**). Given the increased frequency of CD14+ events at the MFI, we asked if these cells express ligands for T cell co-receptors that we found on CD8+ TRM and cytokine-stimulated CD8+ Tmem. We found that mPLAC CD14+ macrophages/monocytes expressed cognate ligands for both 4-1BB (4-1BBL) and PD-1 (PD-L1) on CD8+ TRM (**Figure 4B, C**). Within mPLAC CD14+ events, we observed a subpopulation which expresses high levels of CD206 (**Figure 4C**), phenotypically aligning with descriptions of maternal-derived decidual macrophages(59). In agreement with this definition, CD206hi CD14+ cells were abundant in decidual aspirates but absent from IVB (**Supplemental figure 4B**); further, these CD206hi cells phenocopied the low CD45, CD11b, and CD11c expression also reported in decidual macrophages(59) (**Figure 4D, Supplemental figure 4C**) and did not express fetal sex transcripts (data not shown). Of the CD14+ subsets at the MFI, CD206hi decidual macrophages were the primary producers of 4-1BBL and PD-L1 (**Figure 4D**). Given the high expression of 4-1BBL and PD-L1 on CD206hi decidual macrophages, we asked if these macrophages could physically engage with CD8+ T cells in situ, to provide 4-1BB co-stimulation for tissue residence. While we were unable to leverage CD206 staining in immunofluorescence assays, we used CD163 as a surrogate marker, owing to its high expression on CD206hi decidual macrophages (**Figure 4D**). We found CD45+ leukocytes in distinct aggregates in the decidua basalis; and within these aggregates, we observed CD163+ CD45lo decidual macrophages forming contacts with CD8+ and non CD8+ infiltrates. This indicates that 4-1BB–4-1BBL interactions can occur at the MFI as a possible retention signal (**Figure 4E**).

**Figure 4.**
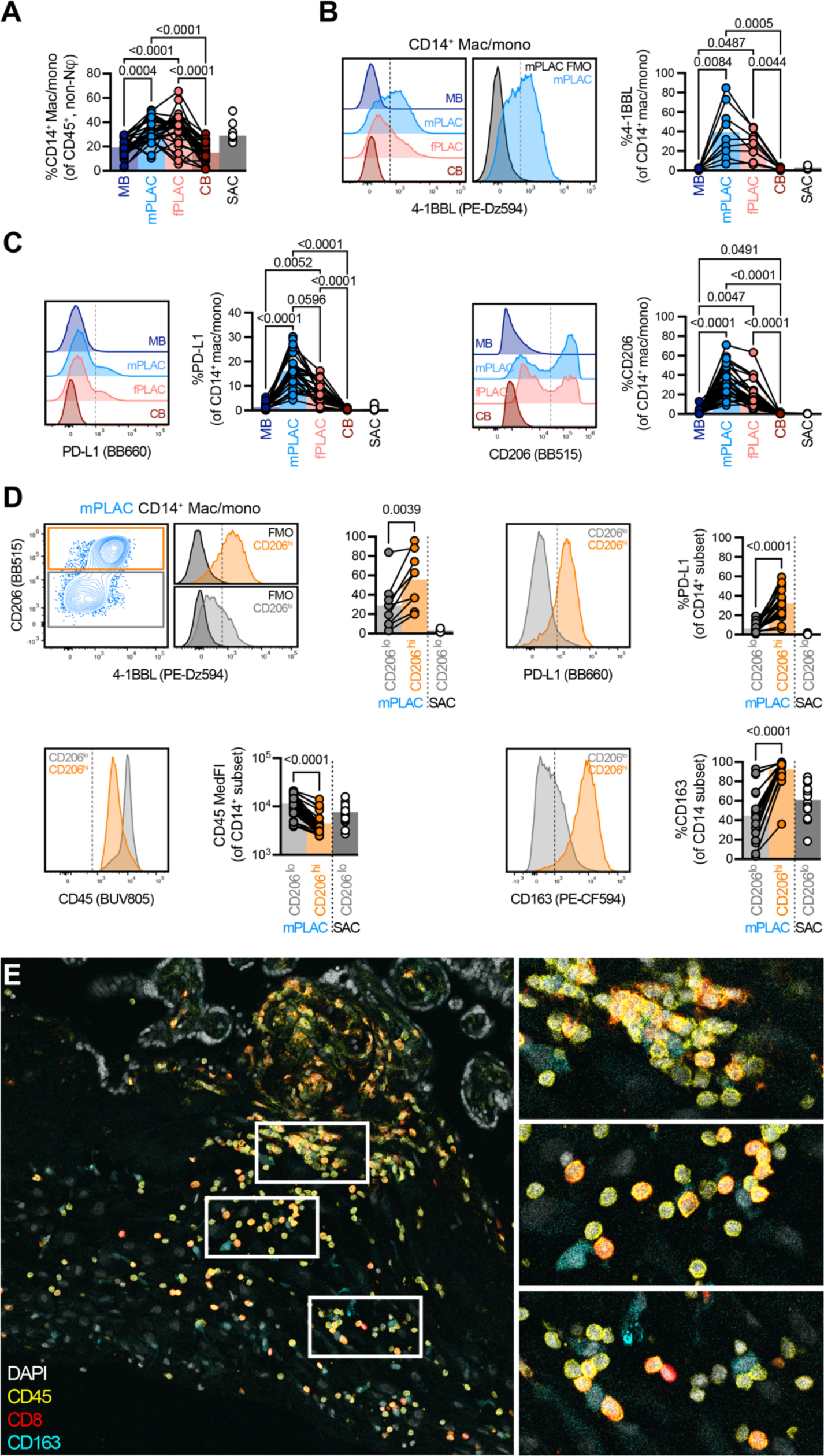
Macrophages at the MFI can engage inflammation-induced receptors on tissue infiltrating T cells. **A** Frequency of CD14^+^ macrophages and monocytes across tissues. **B** 4-1BBL expression within CD14^+^ events across tissues. **C** Expression of PD-L1 and CD206 in CD14^+^ events across tissues. **D** Phenotypic differences of CD206^hi^ and CD206^lo^ CD14^+^ macrophages and monocytes from mfPLAC. **E** Immunofluorescence microscopy of CD8^+^ T cells and CD163^+^ cells at the maternal side of the MFI (decidua basalis). **A** depicts *n* = 29 dyads. **B** depicts *n* = 9 dyads. **C** depicts *n* = 26–33 dyads. **D** depicts *n* = 9–31 dyads. Indicated are statistical significances where *p* < 0.1 by **A**–**C** Friedman tests with Dunn’s multiple comparisons tests or **D** Wilcoxon tests. Symbols in **A**– **D** are connected by donor identity.

### Cytokine and metabolite networks mitigate inflammation-mediated effector programs without compromising cellular activation

Although inflammation upregulates CD8+ Tmem expression of co-stimulatory receptors involved in tissue residence, it also elicits a cytotoxic effector program, termed bystander activation(22). Indeed, our in vitro stimulations with IL-6, IL-15, and IL-18 in combination were able to elicit IFNγ and GzmB upregulation (**Supplemental figure 5A**); yet IFNγ levels at the MFI in plasma or tissue lysates remained undetectable (**Figure 5A, Supplemental figure 5B**). We therefore hypothesized that regulatory mechanisms may be present at the MFI to restrain cytotoxicity elicited by pro-inflammatory signals. We chose to test IL-10, given its role in Treg-mediated control of T cell functions, and found limited levels across all plasma and tissue samples (**Figure 5A, Supplemental figure 5B**). We also surveyed TGF-β1 levels using ELISA, given its presence in NicheNet analysis (**Figure 3B**), and found significantly higher levels in IVB plasma (**Figure 5A**). Although TGF-β1 levels were highest in IVB plasma (**Figure 5A**), detection was impaired in tissue lysate supernatants potentially reflecting limitations in directly comparing tissues and plasmas (**Supplemental figure 5B**). Given the role of L-tryptophan (Trp) metabolism in pregnancy(60) and immune cell fate(61–63), we screened its presence across the depth of the MFI using liquid chromatography-mass spectrometry (LC-MS). Complementing the bias of immunoregulatory cytokines, we found elevated levels of the Trp metabolite, L-kynurenine (KYN), at the maternal versus the fetal side of the MFI (**Figure 5B**). We excluded that MB perfusing the placenta drove this bias, as MB demonstrated significantly lower KYN levels in comparison to CB (**Supplemental figure 5C**). Since KYN must bind to cytosolic aryl hydrocarbon receptor (AhR) to induce AhR nuclear translocation and gene expression(64), we asked if mPLAC CD8+ TRM could take up KYN. We stained for the large neutral amino acid transporter molecule, CD98, responsible for tryptophan and tryptophan metabolite uptake(65). We found mPLAC CD8+ TRM expressed higher amounts of CD98 than their circulating counterparts (**Figure 5C**). Since PD-1 intensity in CD8+ TRM correlated with higher CD98 staining (**Figure 5C**), we asked if cytokine-mediated activation could underlie this expression pattern. Using in vitro T cell stimulations, we found that durable exposure to IL-6, IL-15, and IL-18 could substantially raise CD98 expression (**Figure 5D**). Together, these data suggest TGF-β1 and KYN networks can be sensed by CD8+ TRM at the maternal side of the MFI.

**Figure 5.**
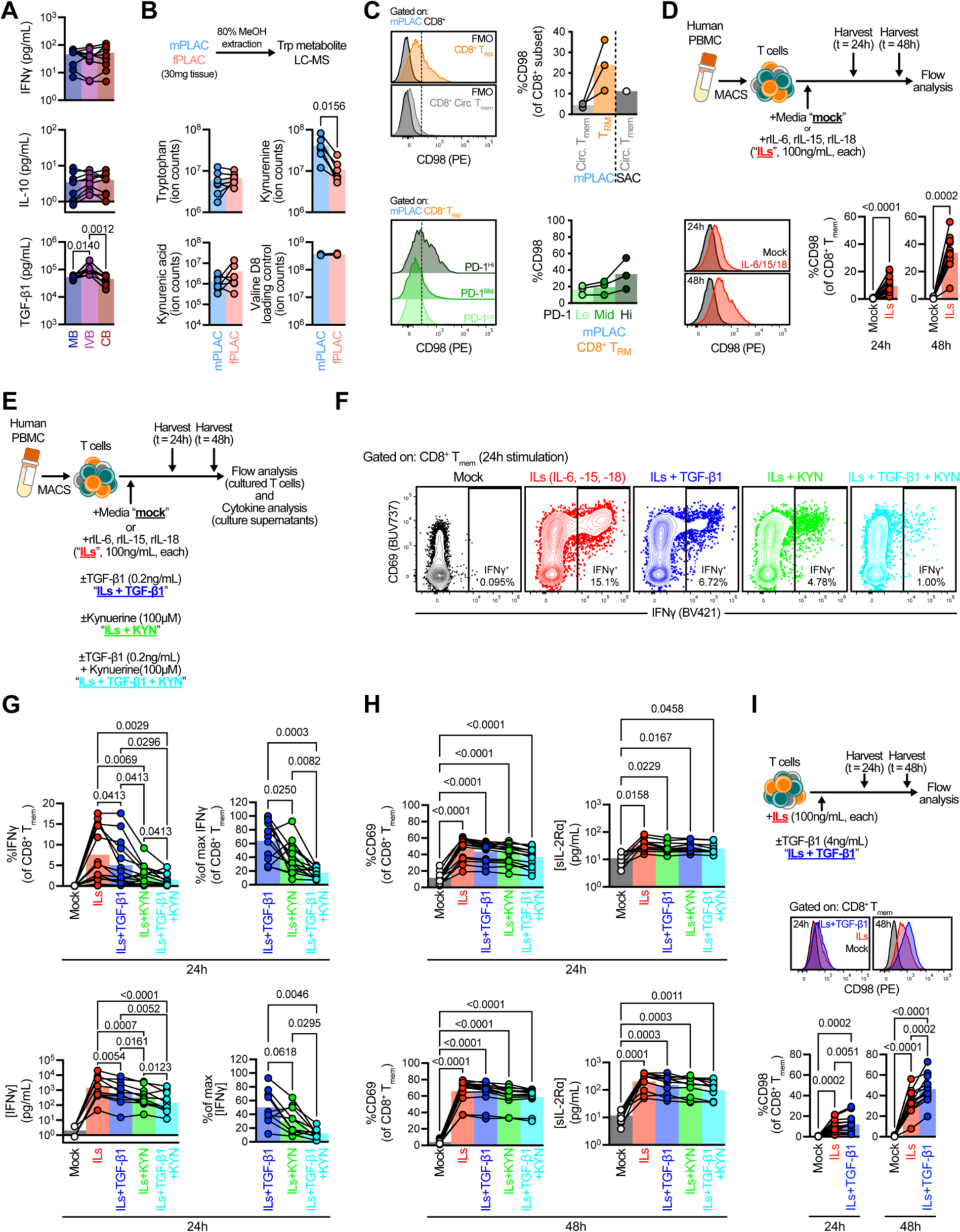
Inhibitory cytokine and metabolite networks at the MFI cooperatively limit inflammation-dependent effector function in CD8^+^ T_mem_ **A** IFNγ, IL-10, and TGF-β1 concentrations across MB, IVB, and CB plasma. **B** LC-MS analysis of tryptophan metabolites and loading controls in mfPLAC and fPLAC lysates. **C** Expression of tryptophan/kynurenine transporter, CD98, in CD8^+^ T_RM_ and circulating T_mem_ from mfPLAC (top) and across PD-1 intensities in mfPLAC CD8^+^ T_RM_. **D** CD98 expression in CD8^+^ T_mem_ after in vitro stimulation with IL-6, IL-15, and IL-18 (100ng/mL, each). **E** Kynurenine and TGF-β1 stimulation assay overview, in which T cells were MACS-isolated from PBMC, cultured with IL-6, IL-15, and IL-18 (100ng/mL, each) in the presence or absence of TGF-β1 (0.2ng/mL) and/or kynurenine (KYN, 100µM), and subsequently analyzed using flow cytometry and multiplex analysis. **F** Representative flow plots of IFNγ expression in CD8^+^ T_mem_ across stimulation conditions after 24h. **G** IFNγ expression in CD8^+^ T_mem_ (left) and IFNγ concentration in culture supernatant (right) across stimulation conditions after 24h. **H** Activation status via CD69 expression in CD8^+^ T_mem_ (left) and sIL-2Rα concentration in culture supernatant (right) across stimulation conditions after 24 and 48h. **I** CD98 expression after stimulation with IL-6, IL-15, and IL-18 (“ILs”; 100ng/mL, each) in the presence or absence of TGF-β1 (4ng/mL). **A** depicts *n* = 9 dyads. **B** depict *n* = 7 placentas. **C** depicts *n* = 3 placentas. **D** depicts *n* = 13–15 PBMC donors. **G–I** depict *n* = 13–15 and *n* = 10 PBMC donors analyzed via flow and multiplex cytokine analysis, respectively. Symbols are connected by donor identity. Indicated are statistical significance where *p* < 0.1 by **A** Friedman tests with Dunn’s multiple comparisons tests, **D** Wilcoxon tests, **G–I** RM one-way ANOVA with Geisser-Greenhouse correction and Holm-Šídák multiple comparisons tests against **G, I** all experimental columns or **H** mock-treated columns.

We next tested whether TGF-β1 and KYN could inhibit CD8+ Tmem effector functions caused by inflammation. We isolated T cells from PBMC with MACS and stimulated them with IL-6, IL-15, and IL-18 in combination (each at 100ng/mL) in the presence of TGF-β1 (at 0.2ng/mL) and/or KYN (at 100µM). To interrogate cytokine expression on a per-cell basis, we added Golgi inhibitors 6 hours prior to harvesting cells and conducting stains for surface expression of activation markers and intracellular cytokine accumulation (**Figure 5E**). Though TGF-β1 or KYN alone could limit inflammation-mediated IFNγ expression in CD8+ Tmem, combinations of both demonstrated cooperativity in attenuating this specific effector function (**Figure 5F, G**). We observed similar cooperativity between TGF-β1 and KYN in T cell culture supernatants in limiting the overall concentrations of IFNγ (**Figure 5G, H**). While TGF-β1 and KYN profoundly reduced IFNγ expression caused by pro-inflammatory signals, its effects were less pronounced on other effector molecules induced by inflammation, like GzmB (**Supplemental figure 5D–E**). Given the limited effects of TGF-β1 and KYN on the upregulation of GzmB, we hypothesized that TGF-β1 and KYN selectively limit IFNγ expression and secretion, rather than impair inflammation-mediated activation in its entirety. Therefore, we tested for markers of cellular activation, CD69 on CD8+ Tmem and soluble IL-2Rα (sIL-2Rα) in culture supernatants. While stimulation with IL-6, IL-15, and IL-18 was a potent inducer of CD69 and sIL-2Rα, TGF-β1 and/or KYN were unable to abrogate their expression (**Figure 5H**). Since TGF-β1 and/or KYN were unable to abolish cytokine-mediated CD8+ Tmem activation, we tested whether these cells also maintained expression of receptors associated with tissue residence. Though TGF-β1 and KYN were able to impair the overall expression of 4-1BB, ICOS, or TIM-3 caused by IL-6, IL-15, and IL-18 stimulation, CD8+ Tmem still expressed these receptors at levels higher than baseline (**Supplemental Figure 6**). Therefore, cellular activation that could beget tissue residence is not entirely forfeited by TGF-β1 and KYN networks.

Since TGF-β1 and KYN inhibited IFNγ to a greater degree in unison, rather than alone, we sought mechanisms that could underlie this cooperativity. We previously found that IL-6, IL-15, and IL-18 could together induce the expression of the KYN transporter, CD98 (**Figure 5D**); however, it was unclear whether TGF-β1 could modify CD98 expression. Using in vitro stimulations of MACS-isolated T cells, we found that the addition of TGF-β1 (at 4ng/mL) to IL-6, IL-15, and IL-18 boosted CD98 expression in CD8+ Tmem by both frequency and median fluorescence intensity (MedFI) (**Figure 5I**). This phenomenon was most apparent at the 48h timepoint, suggesting that cells durably exposed to these signals (like those retained at the MFI) may change transporter expression to properly adapt to their environment.

### Cytokine and metabolite networks selectively permit TCR-mediated effector functions

TCR ligation can also induce both the upregulation of CD98 and IFNγ in CD8+ Tmem. Therefore, we asked if TGF-β1 and KYN cooperatively limits IFNγ expression across T cell activation mechanisms (i.e., both inflammation- and TCR-mediated) or if these molecules solely attenuate IFNγ levels during bystander activation. To test this, we stimulated T cells MACS isolated from PBMC samples with IL-6, IL-15, and IL-18 (each at 100ng/mL) or TCR-agonizing anti-CD3/CD28 microbeads (at a 1:1 bead:cell ratio) in the presence or absence of TGF-β1 (0.2ng/mL) and KYN (100µM) (**Figure 6**). Afterwards, we determined IFNγ concentrations in culture supernatants and interrogated IFNγ expression in CD8+ Tmem using flow. When we stimulated T cells with IL-6, IL-15, and IL-18 in the presence of TGF-β1and KYN, we observed secreted IFNγ levels decrease by a full order of magnitude; however, TGF-β1 and KYN had negligible effect on IFNγ induced by TCR ligation (**Figure 6**). Despite TGF-β1 and KYN failing to attenuate IFNγ expression in TCR-mediated CD8+ Tmem, we observed that TGF-β1 and KYN could still be sensed by these T cells. Specifically, bulk T cell responses (secreted IL-2 and TNFα) and outward signs of activation in CD8+ Tmem elicited by TCR stimulation were slightly impaired by TGF-β1 and KYN (**Supplemental figure 7A–C**). Together, this suggests that while TGF-β1 and KYN can be sensed by CD8+ Tmem across all activation states, that their inhibitory effect on IFNγ expression is most focused to CD8+ Tmem activated by pro-inflammatory cytokines.

**Figure 6.**
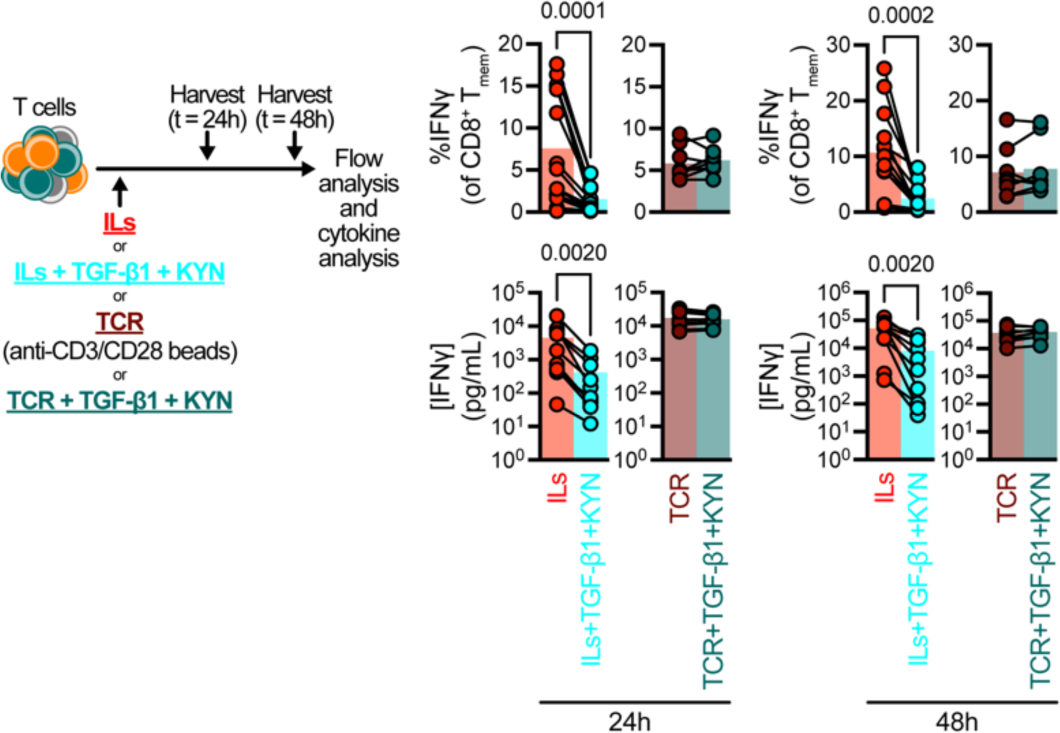
TGF-β1 and kynurenine cooperation selectively impairs cytokine-mediated, but not TCR-mediated, effector responses. IFNγ expression in CD8^+^ T_mem_ and culture supernatant after IL-6, IL-15, and IL-18 (“ILs”; 100ng/mL, each) or anti-CD3/CD28 microbead (“TCR”; 1:1 bead:cell ratio) stimulation in the presence or absence of TGF-β1 (0.2ng/mL) and kynurenine (100µM) after 24 and 48h of culture. Bar plots depict *n* = 8–15 PBMC donors. Symbols in plots are connected by donor identity. Indicated are statistical significance where *p* < 0.1 by Wilcoxon tests.

## Discussion

Maternal-fetal immune conflict during pregnancy has been previously understood as an act of avoidance. In these models, maternal immune cells fail to detect fetal elements because they are denied access to both the MFI (7) and the Ag presentation molecules necessary for TCR-mediated activation and target killing (3). We initially asked if T cells at the term, unlabored MFI mirror those in circulation, in line with the placenta as an “immunologically silent” organ. Contrasting with Nancy, et al. (7) and consistent with recent work from the Tilburgs lab (18), we observed a sizable immune infiltrate largely in the decidua basalis (mPLAC). Using single-cell targeted proteomics and untargeted transcriptomics, we found that a subset of maternal CD8+ T cells at the MFI exhibited a tissue-resident phenotype, which was corroborated by their absence in intervillous circulation. Although Ag presentation is impaired at the MFI(3), we observed a paradoxical signature conventionally associated with TCR activation in these CD8+ TRM, including 4-1BB, PD-1, TOX, and CD39 expression. Therefore, we conducted tetramer screens to identify previously defined Y-encoded antigens (PCDH11Y, USP9Y, DDX3Y, UTY, SMCY/JARID1D) (17). Though we were unable to identify Y Ag-specific CD8+ T cells in MB or mPLAC, we found a wide array of T cells specific for viral Ags. Given the role of cognate Ag and TCR signals in the establishment of tissue residence across multiple organs (36, 66–70), we asked if CD8+ TRM were only specific for chronic viruses, which could reactivate and provide Ag (71). Instead, we also detected CD8+ TRM specific for Ag from acute viral infections, like influenza A virus (IAV) and adenovirus (AdV). These bystander CD8+ Tmem also expressed markers of activation. Though we cannot formally exclude the possibility of an alloresponse leading to TCR agonism (52), TCR engagement should promote T cell proliferation and enhanced numbers in tissues. Rather, we observed that virus-specific CD8+ Tmem formed a similar percentage of the circulating and resident CD8+ Tmem pool, inconsistent with expansion. Together, these data support a model where maternal T cells are retained at the MFI in an Ag-agnostic manner.

Using an untargeted transcriptomic approach, we extricated signatures of chemokine-mediated T cell recruitment (CXCL10–CXCR3) and inflammation-dependent T cell activation (IL15– IL2RG, IL2RB), which we validated with targeted cytokine analyses. The presence of factors capable of recruiting T cells to the MFI contrasts with a previous study describing T cell exclusion from the MFI by means of epigenetically silencing *Cxcl10* (7). Instead, this finding complements other mechanistic studies which found CXCR3-mediated recruitment to the skin (36) and vagina (68) antecedent to residence. As surface CXCR3 levels drop after ligand engagement (55, 72), the decreased CXCR3 frequency in circulating CD8+ Tmem at the MFI may reflect CXCR3-CXCL10 signaling in action. But this recruitment to the MFI ultimately exposes CD8+ Tmem to an entirely different environment, in which there is sustained exposure to pro-inflammatory cytokines, like IL-6, IL-15, and IL-18. Using in vitro stimulations, we found that these pro-inflammatory signals could recapitulate the phenotypes observed in mPLAC CD8+ TRM, including PD-1, 4-1BB, ICOS, TIM-3, TOX, and CD39 expression. CD39 has been proposed as a marker of recent TCR activation; but recent work identified paradoxical CD39 expression in decidual T cells with TCRs specific for irrelevant viral antigens (18). Our study identifies a cytokine-dependent, TCR-independent mechanism to CD39 expression that may explain this phenomenon, adding to a growing number of studies elucidating parallels in the outcomes of inflammation- and TCR-mediated activation in CD8+ Tmem (22, 73). These similarities likely explain why CD8+ TRM paradoxically upregulate transcripts associated with TCR engagement despite the probable lack of cognate Ag: homeostatic pro-inflammatory cytokine networks within tissues promote these phenotypic adaptations.

Ag-nonspecific CD8+ TRM are not often described in mice, likely because of the limited CD8+ Tmem pool in animals reared in SPF settings (74). Given this limitation, signals necessary for tissue residence have been most understood downstream of TCR activation. 1) TCR ligation induces 4-1BB expression in CD8+ T cells and 2) subsequent 4-1BB–4-1BBL signaling is required for subsequent CD8+ T cell tissue residence (57, 58). However, our broad analysis of the placenta demonstrates that inflammation may serve as a substitute for TCR agonism: IL-15 maintains 4-1BB expression in CD8+ Tmem despite the lack of cognate Ag, which can be engaged in situ by 4-1BBL–expressing CD206hi macrophages. Though 4-1BB signals are critical for residency in select tissues (66, 67), we also show other receptors important to residency, like ICOS (75), can be upregulated by inflammation. Though outside the scope of this current study and limited in availability, incorporating tissues from donors on immunosuppressants that limit TCR-mediated activation (76) could provide further granularity regarding the signals that drive residency at the MFI.

Given the likely importance of pro-inflammatory cytokines in tissue retention, but their paradoxical role in bystander activation (22), we sought mechanisms which could selectively impair inflammation-driven cytotoxicity in CD8+ Tmem. Using comprehensive transcriptomics, proteomics, and metabolomics, we identified that the pro-inflammatory cytokine milieu at the MFI exists in juxtaposition with anti-inflammatory cytokine (TGF-β1) and metabolite (KYN) networks. IL-6, IL-15, and IL-18 can activate CD8+ Tmem and lead to the acquisition of effector functions, like IFNγ secretion, which could threaten immune homeostasis; however, TGF-β1 and KYN can cooperatively restrain inflammation-mediated IFNγ expression without abrogating cellular activation and coreceptor upregulation. Finally, we do not observe these effects on cytotoxicity with TCR-stimulated T cells, suggesting this regulatory mechanism serves two purposes: 1) to broadly sustain CD8+ TRM at the MFI in situ without inducing potentially detrimental inflammation-mediated cytotoxicity and 2) to concurrently maintain maternal immunocompetence against pathogens by selectively permitting TCR-mediated cytotoxicity. Incorporating these regulatory signals appears essential for CD8+ TRM, given their biased expression of the KYN transporter, CD98, alongside reports demonstrating that TGF-β1(36, 67) and AhR(77) are required for their persistence at other organs.

Despite existing at the MFI, these regulatory programs are likely involved in the maintenance of immune homeostasis at other tissues characterized by pro-inflammatory microenvironments. Given the role of dysregulated inflammation (often IL-15–related) and bystander activation in various autoimmune pathologies(23, 78), TGF-β1 and/or KYN-AhR signaling could be leveraged to restore homeostasis in multiple contexts, such as reducing inflammation-associated preterm birth. Conversely, leveraging these programs may have therapeutic relevance in cancer, as tumors are often enriched for TGF-β1(79) and KYN (63). But it remains unclear these regulatory molecules could be targeted so that tumor CD8+ TRM could be activated into killers by inflammation alone. Future work will be necessary to dissect the roles of TGF-β1, KYN, and pro-inflammatory cytokines across tissue types and disease states to better control CD8+ Tmem functions.

In summary, we propose that Ag-nonspecific CD8+ Tmem recruitment and retention can occur via pro-inflammatory homeostatic cytokine networks in situ; but immune homeostasis is maintained because inflammation-mediated cytotoxicity programs are restrained by contemporaneous sensing of regulatory cytokines and metabolites.

## Methods

### Study population and approvals

We obtained samples from pregnant individuals at the University of Washington Medical Center (Seattle, WA) under Institutional Review Board (IRB) approved studies from the University of Washington (STUDY00001636). All individual participants gave written informed consent prior to participation. We included pregnant individuals >18 years old with non-anomalous singleton fetus undergoing a scheduled cesarean delivery at term gestation (between 37-41 weeks gestation). Individuals with pregnancy complications including multiple gestation, pre- or gestational diabetes, preeclampsia, preterm labor, or placental abnormalities were excluded. HIV-uninfected adults were recruited by the Seattle HIV Vaccine Trials Unit (Seattle, Washington, USA) as part of the study “Establishing Immunologic Assays for Determining HIV-1 Prevention and Control.” These samples are also known as the Seattle Area Control (SAC) cohort. All participants were provided and signed informed consent, and the Fred Hutchinson Cancer Research Center Institutional Review Board approved the study protocol.

### Blood collection for leukocyte and plasma isolation

We maintained all samples on ice after acquisition. Clinicians collected maternal peripheral blood (MB) prior to delivery and fetal cord blood (CB) immediately after delivery by venipuncture. MB and CB were collected in acid citrate dextrose solution A (ACD)-vacutainer tubes (BD Biosciences). To collect intervillous blood (IVB), we injected 6–10mL of ACD solution using a 28G½ syringe into the intervillous space of the placenta through the fetal membranes to prevent thrombosis (**Supplemental figure 2C**). We then obtained IVB which passively drained from the placenta and processed it alongside MB and CB samples. We centrifuged MB, CB, and IVB at 400g for 10 min, collected plasma and stored at -80°C. We isolated leukocytes via ACK lysis (ThermoFisher) or density gradient centrifugation with Lymphoprep (STEMCELL Technologies) and resuspended in RP7.5 (RPMI 1640 supplemented with 7.5% FBS (Peak Serum), 2 mM L-glutamine, 100 U/mL penicillin–streptomycin (ThermoFisher)) before downstream use. To quantitate cells, we stained single-cell suspension aliquots (**Supplemental table 1**) and acquired events on a Guava easyCyte (EMD Millipore). We then stained cells immediately for flow analysis or cryopreserved cells in cryopreservation solution (CPS; 10% sterile DMSO (Sigma Aldrich), 90% FBS).

### Tissue collection for leukocyte isolation

We maintained all tissues on ice after acquisition. Clinicians vacuum aspirated residual tissue in the uterine cavity to obtain decidual tissue. We rinsed intact placentas in 1x PBS and used sterile instruments to collect 0.5cm-depth biopsies from the maternal-facing and fetal-facing sides of the placenta (mfPLAC and fPLAC, respectively). We minced decidual, mfPLAC, and fPLAC tissues in digestion media: RP7.5 + 700U/mL Collagenase Type II (Sigma Aldrich), 200U/mL DNase (Sigma Aldrich), which we incubated in a shaking 37°C incubator for 60 min at 225 RPM. Afterwards, we strained single-cell suspensions through a 70µm filter, centrifuged cells at 400g for 5 min, discarded supernatants, and washed cells with RP7.5 media. We removed RBCs using ACK lysis, and resuspended cell pellets in RP7.5 before downstream use. To quantitate cells, we stained single-cell suspension aliquots (**Supplemental table 1**) and acquired events on a Guava easyCyte. We then stained cells immediately for flow analysis or cryopreserved cells in cryopreservation solution (CPS; 10% sterile DMSO (Sigma Aldrich), 90% FBS).

### In vitro stimulation assays

We thawed 2–4 × 10^7^ cryopreserved PBMC in RP10 media (RPMI1640 supplemented with 10% FBS, 2 mM L-glutamine, 100 U/mL penicillin-streptomycin). To enrich bulk T cells from single-cell suspensions, we used human-specific T cell negative isolation MACS (STEMCELL Technologies). We plated 0.5–2 × 10^6^ T cells per well in 96-well V-bottom tissue culture plates. We cultured cells in RP10 at 37°C and 5% CO_2_. We stimulated T cells with human recombinant IL-6 (BioLegend), IL-15, and IL-18 (Peprotech) in combination (each at 100ng/mL) or with Dynabeads human T-Activator (ThermoFisher) anti-CD3/CD28 beads (at a 1:1 bead/cell ratio). We added 100µM L-kynurenine (Sigma) dissolved in RP10 and/or human recombinant TGF-β1 (Peprotech) at the time of stimulation. Six hours prior to harvesting T cells, we added GolgiPlug (BD Biosciences) at 1:1000 dilution to allow intracellular accumulation of cytokines. At harvest timepoints (*t* = 24 or 48h), we centrifuged plates, collected and froze media supernatants at -80°C, and conducted flow cytometric staining on cells.

### Flow cytometry

When handling cryopreserved samples for ex vivo cytometric analysis, we thawed single-cell suspensions in warm RP10 (RPMI 1640 supplemented with 10% FBS, 2 mM L-glutamine, 100 U/mL penicillin–streptomycin). We aliquoted fresh or thawed single-cell suspension samples at 0.5–2 × 10^6^ cells/well in a 96-well v-bottom plate. All flow panel reagent information, stain conditions, and gating strategies are included in **Supplemental figures 8–14, Supplemental tables 2–8**. We conducted LIVE/DEAD fixable aqua or blue viability dye (Thermo Fisher) (AViD or BViD, respectively) staining in 1x PBS. For FACS surface staining, we utilized FACSWash (1x PBS supplemented with 2% PBS) as the stain diluent; and for all other flow analysis, we utilized FACS Wash + Azide (1 × PBS supplemented with 2% FBS and 0.2% sodium azide) as the stain diluent. For T cell panels (**Supplemental table 2, 4, 5, 6**) we fixed cells with the FOXP3 Fixation/Permeabilization Buffer Kit (Thermo Fisher) and conducted intracellular/nuclear stains using the FOXP3 Permeabilization Buffer (Thermo Fisher) as diluent. For APC panels (**Supplemental tables 7, 8**) we fixed cells with Cytofix/Cytoperm (BD Biosciences) and conducted intracellular stains using 1x Perm/Wash buffer (BD Biosciences) as diluent. We resuspended cells in FACS Wash and acquired events on a FACSymphony A5 and sorted cells on a FACSAria II (BD Biosciences), which we analyzed using FlowJo v10 (BD Biosciences). We conducted statistical testing using Prism v8 (GraphPad).

### CyTOF tetramer screening

The Fred Hutchinson Cancer Research Center Immune Monitoring Core refolded HLA-A*01:01, HLA-A*02:01, HLA-A*03:01, HLA-A*11:01, and HLA-B*07:02 monomers with appropriate UV-cleavable peptides and biotinylated them. We labeled streptavidin (SA) with heavy metals as previously described(50). We purchased purified antibodies lacking carrier proteins, which we conjugated to heavy metals using DN3 polymers and according to protocols provided by Fluidigm.

We conducted multiplex tetramers staining in accordance with previously published protocols(80). Briefly, we labeled each tetramer with a combination of two different metal-labeled SAs. Using ten different SA conjugates (**Supplemental table 9**), we generated 45 possible combinations. We assigned each specific combination to a single peptide (**Supplemental tables 10, 11**). We mixed SAs (50µg/mL, 30µL) in the assigned combination for each peptide in a 96-well plate. In another 96-well plate, we added 5µL of peptide (1mM) to 100µL of HLA monomer (0.1mg/mL, diluted in PBS), with a different peptide in each well. We exposed peptide-MHC plates to UV light for 10 min and then incubated at 4°C overnight. After incubating, we tetramerized SAs (50µg/mL) to the assigned peptide-MHC complex by adding 3 additions of 10µL of each metal-labeled double-coded SA into peptide-MHC wells (3x 10µL) according to our coding scheme, with 10 min incubations between each addition. We then incubated tetramerized peptide-MHC complexes with 10µM free biotin (Avidity) for 10 min. We then combined all tetramers and concentrated in a 50-kDa Amicon filter (Millipore) to a final volume of 500µL. We added designated antibodies (1° surface stain, see **Supplemental table 12**), and adjusted the final cocktail volume to 1mL with PBS + 0.5% FBS and 0.02% sodium azide. Prior to use, we filtered the tetramer/antibody cocktail with a 0.1µm filter (Millipore).

We thawed cryopreserved MB and mfPLAC samples where maternal HLA types were HLA-A*01:01, HLA-A*02:01, HLA-A*03:01, HLA-A*11:01, and/or HLA-B*07:02 in warm RP10 + 10µg/mL DNase. We stained samples in accordance with previously published protocols (80). Briefly, we stained cells with 1° surface stain cocktail (including tetramers) for 60 min at room temperature. We then conducted viability staining using 12.5µM cisplatin in 1x PBS for 10 min on ice. We prepared 2° surface stain antibody cocktails (**Supplemental table 12**) in advance, which we froze into multiple aliquots to minimize batch variance. We thawed and stained cells with this cocktail for 20 min on ice. Afterwards, we washed and fixed cells overnight in 2% PFA. We washed cells with 1x eBioscience FOXP3 permeabilization buffer, which we also used as a diluent for intracellular antibody staining (30 min at room temperature). Prior to acquisition, we stained cells with Cell ID intercalator Ir (Fluidigm) for 10 min on ice and then washed 3x with Milli-Q H_2_O. We then acquired samples using Helios (Fluidigm), analysed using FlowJo v10, and conducted statistical testing with Prism v8.

### Single-cell library preparation and sequencing

We generated cDNA libraries from sorted cell populations (CD8^+^ T cell and CD45^+^ CD3^-^ CD19^-^ HLA-DR^+^ APCs) using the Chromium Single Cell 3’ Reagent Kits (v2 protocol) or the Single Cell 5’ Reagent Kit (v1 protocol) (10x Genomics). We used the Chromium Controller (10x Genomics) to generate oil emulsion droplets of single cells with barcoded gel beads and reverse transcriptase mix. We generated barcoded cDNA within these droplets, and purified cDNA using DynaBeads MyOne Silane magnetic beads (ThermoFisher). We amplified cDNA by PCR (10 cycles) using reagents from the Chromium Single Cell 3’ Reagent Kit (v2) or the GEX Reagent kit (v1). We purified amplified cDNA with SPRIselect magnetic beads (Beckman Coulter) according to their protocol.We enzymatically fragmented and size-selected cDNA prior to library construction, which we conducted by performing end repair, A-tailing, adaptor ligation, and PCR (12 cycles). In order to assess library quality, we used Agilent 2200 TapeStation with High Sensitivity D5000 ScreenTape (Agilent). We assessed library quantities using digital droplet PCR (ddPCR) with the Library Quantification Kit for Illumina TruSeq (BioRad). We diluted libraries to 2nM and performed paired-end sequencing using a HiSeq 2500 (Illumina).

### Single-cell RNA sequencing analysis

We utilized the R package, Seurat (81), for all downstream analysis, following general guidelines for analyzing scRNAseq data. For our analyses, we only included cells that had ≥ 200 genes, and depending on sample distribution, ≤ 10% mitochondrial genes. We merged all acquired samples into a single Seurat object. We then used a natural log normalization with a scale factor of 10,000, determined variable genes using the vst method, and z-score scaled. We used principal component analysis (PCA) to generate 50 PCs which were used to calculate UMAPs, for which we created graph-based clusters using a resolution between 0.2 and 0.6. For non-biased cell annotation, we applied SingleR (82) and used the expression of typical lineage transcripts to verify the cell label annotation. For all differential gene expression analyses, we utilized the Seurat implementation of MAST (model-based analysis of single-cell transcriptomes) (83) with the number of unique molecular indexes (UMIs) included as a covariate (proxy for cellular detection rate (CDR)) in the model. NicheNet (53) analysis was adapted from the vignette described at https://github.com/saeyslab/nichenetr containing T cells and APCs (described above). We subset these populations to contain only immune sorted from mfPLAC biopsies. We set different CD8^+^ T cell subsets as “receiver” (i.e., resident or circulating memory CD8^+^ T cell clusters) and all myeloid cell cluster (except the mast cell cluster) as “sender” populations. We conducted a DE gene test to find genes enriched in mfPLAC over MB samples. We performed NicheNet analysis based on the vignette to infer receptors, filtered for documented links, and generated a circos plot of the ligand-receptor interactions for the top 20 ligands within the respective cellular populations. Main scripts used for data processing are available at https://github.com/Jami-Erickson.

### Tissue collection for protein isolation

We collected 0.5cm-depth mfPLAC and fPLAC biopsies and homogenized tissues in lysing buffer (1x PBS + 0.05% Tween 20 (ThermoFisher)) at 1g tissue/5mL buffer. We centrifuged samples at 10,000g for 5 min at 4°C, collected lysate supernatants, and stored at -80°C.

### Cytokine analysis

We provided plasma, tissue lysate supernatants, and cell culture supernatants to the Fred Hutchinson Cancer Research Center Immune Monitoring Core for ELISA (TGF-β1) or multiplex cytokine analysis (all other factors). Multiplex measurements were analyzed on a Luminex 200 (Luminex Corporation).

### Immunofluorescence

We collected full-depth (i.e., spanning the maternal- and fetal-facing surfaces) biopsies from the placenta and fixed overnight in Cytofix buffer (BD Biosciences) diluted 1:4 in 1x PBS. We dehydrated fixed tissues in sterile-filtered 20% w/v sucrose (Sigma Aldrich), embedded in Tissue-Tek OCT media (Sakura Finetek), froze in the vapor phase of LN_2_, and stored at -80°C. We cut 8–10 µm sections of tissues using a CM1950 cryostat (Leica), dried slides at room temperature overnight, and then stored at -80°C. On the day of staining, we incubated slides in -20°C acetone for 5 min, dried slides, and outlined tissues with a hydrophobic pap pen. We rehydrated slides in 1x PBS for 5 min, and then incubated slides in blocking buffer (1x TBS + 5% normal human serum) for 30 minutes. We conducted stains as described in **Supplemental table 13** and washed slides with TBS-T (1x TBS + 0.05% Tween-20). We mounted slides in ProLong Gold (ThermoFisher), cured slides overnight, and imaged on a SP8 confocal microscope (Leica) at 20x magnification.

### Metabolomics sample preparation

We prepared a reference library of tryptophan (Trp) metabolites for our liquid chromatography mass spectrometry (LC-MS) analyses by dissolving select metabolites in HPLC grade H_2_O at ∼0.5mM (cinnabarinic acid) or ∼5mM (all other metabolites) (**Supplemental table 14**).

We prepared plasma samples for LC-MS by adding 6µL plasma to a microcentrifuge tube with 200µL ice cold HPLC-grade 80% MeOH with 250µM valine D8 standard as a loading control. We vortexed samples for 10 minutes at 4°C, then centrifuged at 17,000g for 10 min at 4°C. We transferred 100µL of supernatant to a new microfuge tube, with 50µL moved to an LC-MS vial for metabolomics analysis.

We prepared flash-frozen mfPLAC and fPLAC biopsies by weighing out 30mg of frozen tissue, which were transferred to a microcentrifuge tube and maintained in LN_2_. We homogenized tissues within microtubes with a miniature LN_2_-cooled mortar and pestle. To extract metabolites, we added 900µL ice-cold HPLC-grade 80% MeOH with 250µM valine D8 loading control and vortexed samples for 10 min at 4°C. We removed debris by centrifuging tubes at 17,000g for 10 minutes at 4°C, and transferred 700µL of supernatant to a new microcentrifuge tube, with 50µL moved to a LC-MS vial for metabolomics analysis.

### LC-MS metabolomics

We transferred samples into LC-MS vials for measurement by LC-MS. We performed metabolite quantitation using a Q Exactive HF-X Hybrid Quadrupole-Orbitrap Mass Spectrometer equipped with an Ion Max API source and H-ESI II probe, coupled to a Vanquish Flex Binary UHPLC system (ThermoFisher). We completed mass calibrations at a minimum of every 5 days in both the positive and negative polarity modes using LTQ Velos ESI Calibration Solution (Pierce). We chromatographically separated samples by injecting a sample volume of 1µL into a SeQuant ZIC-pHILIC Polymeric column (150 x 2.1mm 5µM, EMD Millipore). We set flow rates to 150µL, autosampler temperature to 10°C, and column temperature to 30°C. Our Mobile Phase A consisted of 20mM ammonium carbonate an 0.1% (v/v) ammonium hydroxide, while Mobile Phase B consisted of 100% acetonitrile. Our samples were gradient eluted (%B) from the column as follows: 0–20 min: linear gradient from 80% to 20% B; 20–24 min: hold at 20% B; 24–24.5 min: linear gradient from 20% to 85% B; 24.5 min–end: hold at 85% B until equilibrated with ten column volumes. We directed Mobile Phase into the ion source with the following parameters: sheath gas = 45, auxiliary gas = 15, sweep gas = 2, spray voltage = 2.9 kV in the negative mode or 3.5 kV in the positive mode, capillary temperature = 300°C, RF level = 40%, auxiliary gas heater temperature = 325°C. We conducted mass detection with a resolution of 240,000 in full scan mode, with an AGC target of 3,000,000 and maximum injection time 250 msec. We detected metabolites over a mass range of 70-1050 *m*/*z* in full scan mode with polarity switching. We quantitated all metabolites using Tracefinder 4.1 (ThermoFisher) referencing our in-house kynurenine pathway metabolite standards libraries using ≤ 5 ppm mass error.

## Acknowledgements

We thank the UWMC Labor and Delivery staff and study participants for their generosity. We thank our research study coordinators. We also thank Fred Hutch Shared Resources for technical assistance and Dr. Alec Wilkens, Dr. Antje Heit, and Kimberly Smythe for helpful discussions. We thank Dr. Stephen Jameson for critical review of the manuscript. This work was supported by NIH grants R21 AI144677 (to R.S.), F31 HD098769 (to N.J.M.), TL1 TR002318 (to N.J.M.), and National Cancer Institute grants F99 CA245735 (to N.J.M.) and R01 CA264646 (to E.W.N). E.W.N. is supported by the Andy Hill Endowment Distinguished Researcher CARE fund. S.A.M. is supported by the Reproductive Scientist Development Program K12HD000849 from the Eunice Kennedy Shriver National Institute of Child Health & Human Development and the Burroughs Wellcome Fund. N.J.M. is a Leslie and Pete Higgins Achievement Rewards for College Scientists Fellow and Dr. Nancy Herrigel-Babienko Memorial Scholar, C.S.D. is a Hartwell Fellow.

