## Supplementary figures and tables for "Converging cytokine and metabolite networks shape asymmetric T cell fate at the term human maternal-fetal interface"

### SUPPLEMENTARY MATERIALS

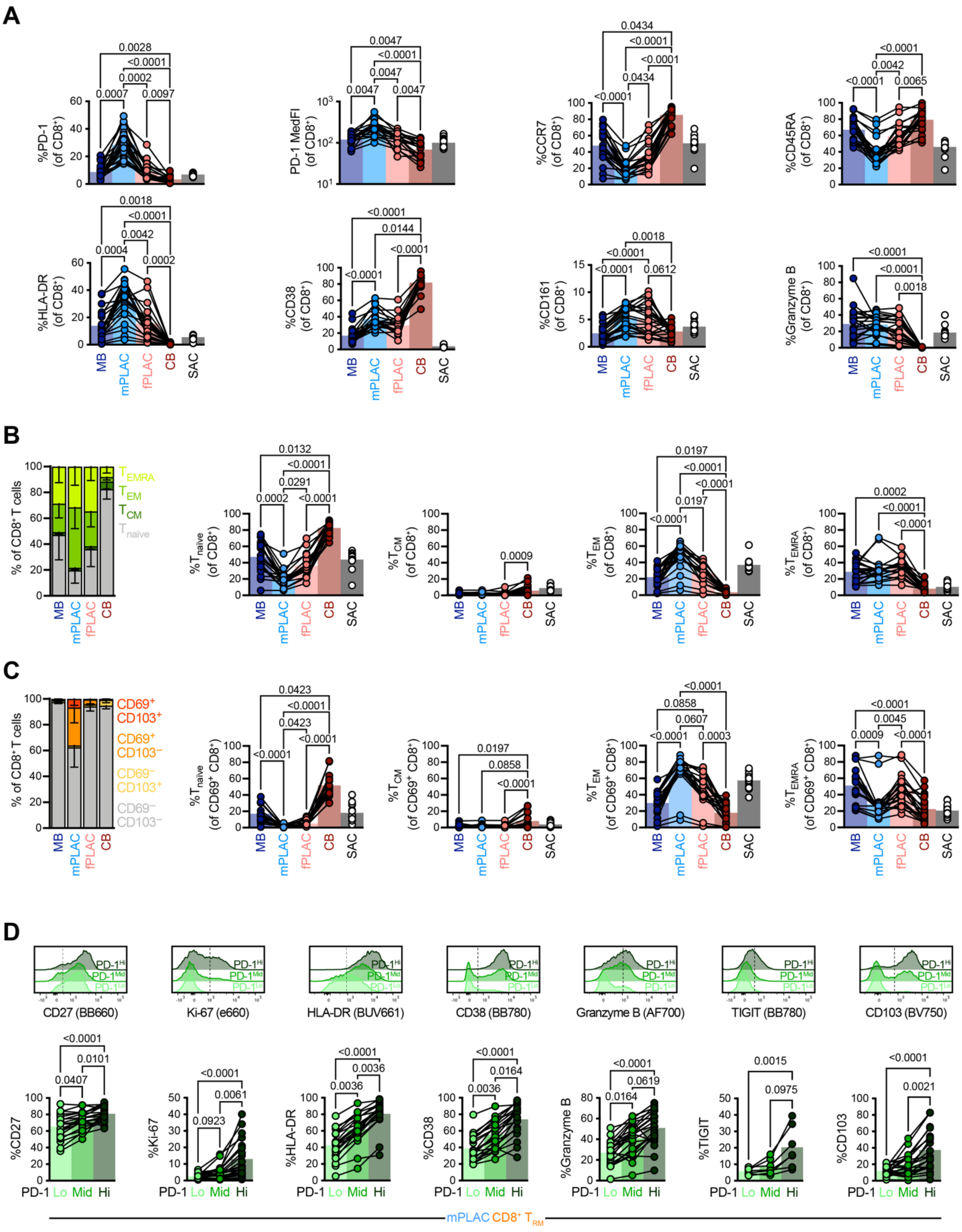
A
%PD-1 (of CD8+)
PD-1 MedFI (of CD8+)
%CCR7 (of CD8+)
%CD45RA (of CD8+)
%HLA-DR (of CD8+)
%CD38 (of CD8+)
%CD161 (of CD8+)
%Granzyme B (of CD8+)
MB
mPLAC
fPLAC
CB
SAC
B
% of CD8+ T cells
TEMRA
TEM
TCM
Tnaïve
%Tnaïve (of CD8+)
%TCM (of CD8+)
%TEM (of CD8+)
%TEMRA (of CD8+)
C
CD69+ CD103+
CD69+ CD103−
CD69− CD103+
CD69− CD103−
%Tnaïve (of CD69+ CD8+)
%TCM (of CD69+ CD8+)
%TEM (of CD69+ CD8+)
%TEMRA (of CD69+ CD8+)
D
PD-1Hi
PD-1Mid
PD-1Lo
CD27 (BB660)
Ki-67 (e660)
HLA-DR (BUV661)
CD38 (BB780)
Granzyme B (AF700)
TIGIT (BB780)
CD103 (BV750)
%CD27
%Ki-67
%HLA-DR
%CD38
%Granzyme B
%TIGIT
%CD103
PD-1 Lo Mid Hi
mPLAC CD8+ TRM

**Supplemental figure 1:**
**$CD8^+$ $T_{RM}$ from the maternal side of the MFI are phenotypically distinct**

**A** Phenotypes of bulk $CD8^+$ T cells across tissue compartments. **B** Memory subset distributions of $CD8^+$ T cells within tissue compartments. **C** Memory subset distributions of $CD69^+$ $CD8^+$ T cell events across tissues. **D** Phenotypic differences across PD-$1^{Hi}$, PD-$1^{Mid}$, and PD-$1^{Lo}$ $CD8^+$ $T_{RM}$ from mPLAC. **A** depicts 13-22 dyads. **B** and **C** depict 20 dyads. **D** depicts 7–21 donors. All points represent a unique population that are connected by donor identity, with bars signifying mean. In bar charts in **B** and **C**, bars depict mean with SD. All statistical significances where $p < 0.1$ are shown as calculated by Friedman tests with Dunn's multiple comparison test.

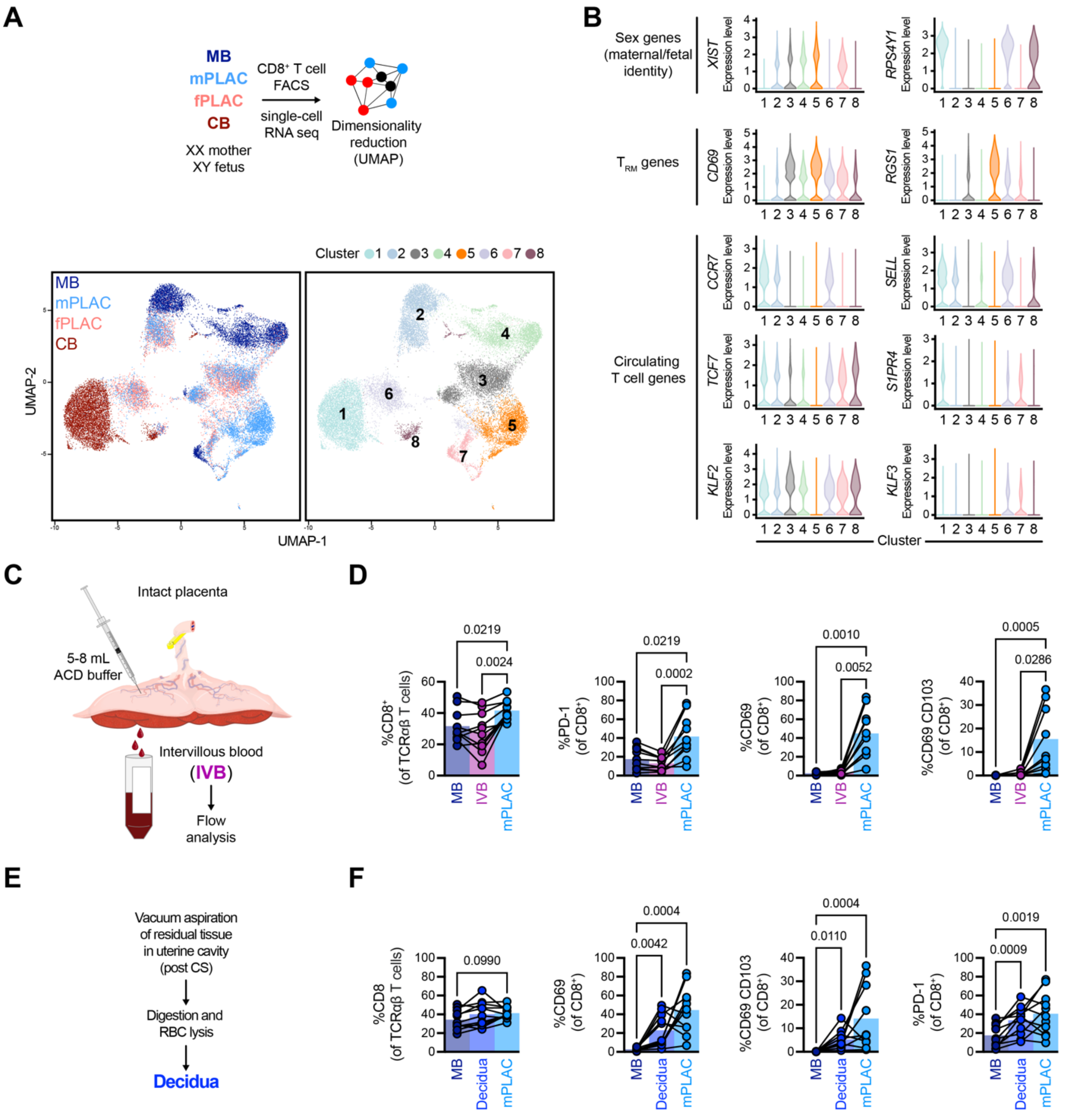


**Supplemental figure 2**
**CD8$^+$ T$_{RM}$ are maternally derived and absent from circulatory elements of the MFI**

**A** Overview of single-cell RNAseq approach, in which CD8$^+$ T cells were isolated from tissues using FACS, and resulting data visualized by UMAP colored by tissue source (left) and graph-based clustering (right). **B** Violin plots of log-transformed normalized transcript counts across clusters for genes indicative of sex (*XIST*, female; *RPS4Y1*, male), tissue residence (*CD69*, *RGS1*), and capacity to circulate (*CCR7*, *SELL*, *TCF7*, *S1PR4*, *KLF2*, *KLF3*). **C** Overview of intervillous blood (IVB) isolation. **D** Frequency of CD8$^+$ T cells and T$_{RM}$-associated proteins within

$CD8^+$ T cell events from MB, IVB, and mPLAC. **E** Overview of decidua collection and processing. **F** Frequency of $CD8^+$ T cells and $T_{RM}$-associated proteins within $CD8^+$ T cell events from MB, decidua, and mPLAC. **A** and **B** depicts 1 dyad with an XY fetus. **D** depicts 9–10 donors. **F** depicts 10–11 donors. All points in **D** and **F** depict a unique population with points connected by donor identity, with bars indicating mean. Statistical significances by Friedman tests with Dunn's multiple comparison tests where $p < 0.1$ are indicated.

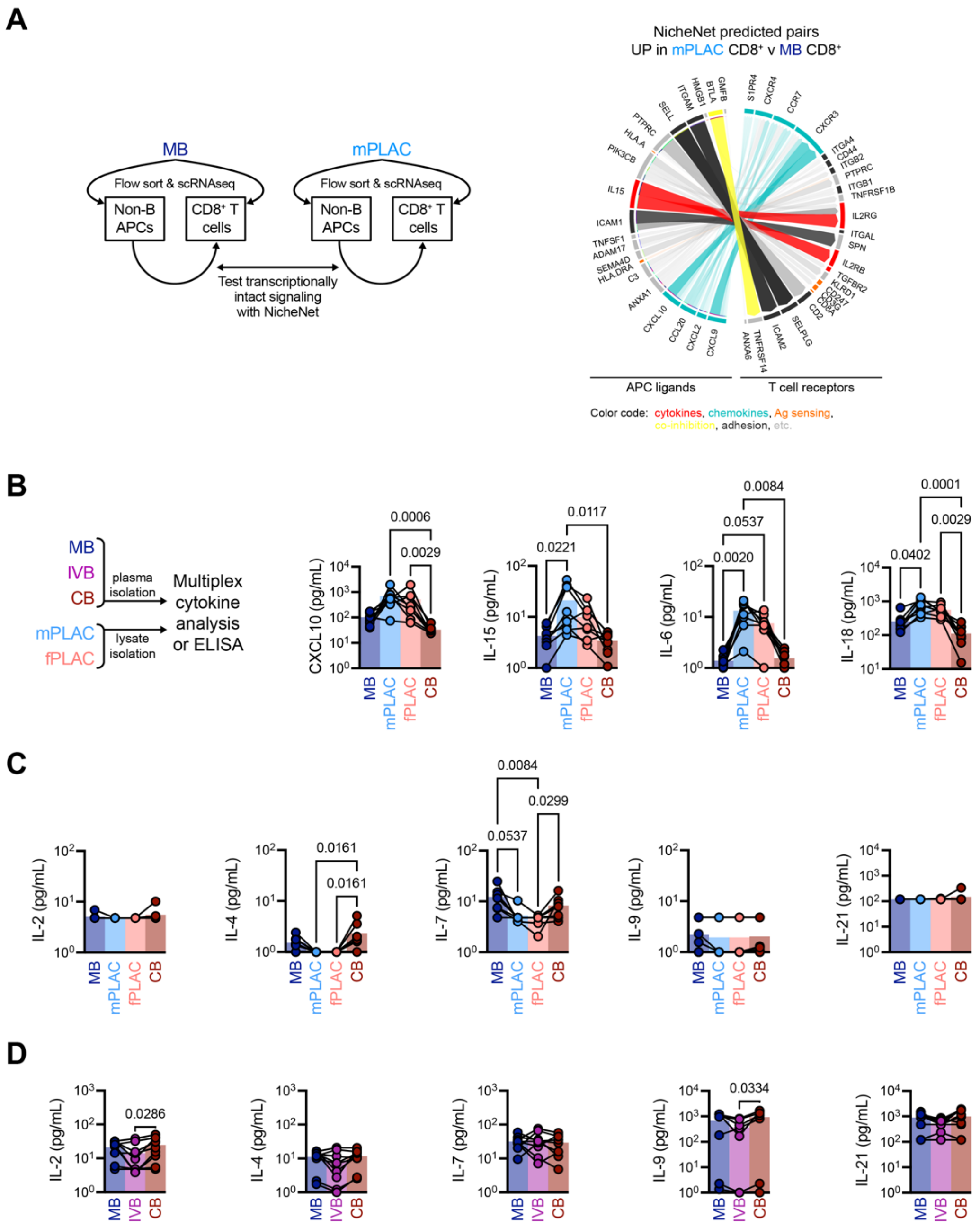


**Supplemental figure 3**
**Other γc cytokines are not elevated at the MFI**

**A** NicheNet analysis of potential ligand-receptor interactions that are enriched in mPLAC APCs and CD8+ T cells versus MB APCs and CD8+ T cells. CIRCOS plot interactions are colored to highlight cytokine (red), chemokine (teal), Ag presentation/sensing (orange), co-inhibitory (yellow), and adhesion (dark grey) interactions enriched in mPLAC APCs and CD8+ T cells over blood. **B** Multiplex analysis of paired MB and CB plasmas, as well as lysates from mPLAC and fPLAC tissues. **C** and **D** Multiplex analysis for $\gamma_c$ cytokines (IL-2, IL-4, IL-7, IL-9, and IL-21) in (**B**) MB and CB plasmas and mPLAC and fPLAC lysates or (**C**) MB, IVB, and CB plasmas. **A** depicts scRNAseq data from *n* = 1 dyad with an XY fetus. **B** and **C** depict *n* = 8 dyads. **D** depicts *n* = 7–9 dyads. Indicated are statistical significances where $p < 0.1$ by Friedman tests with Dunn's multiple comparison tests. Symbols in **B**–**D** are connected by donor identity.

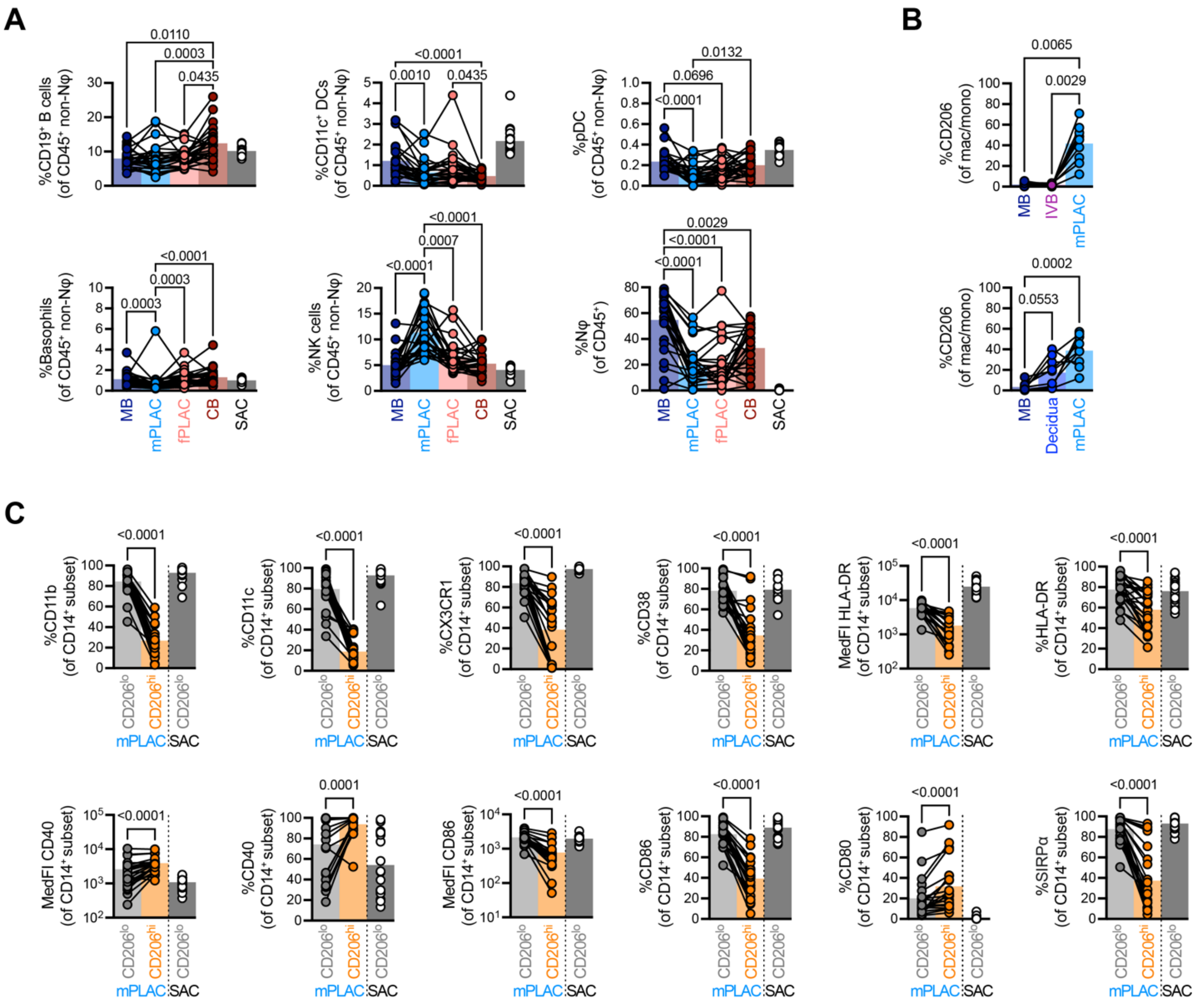


**Supplemental figure 4**
**Phenotypically distinct macrophages occupy the stroma of the MFI**

**A** Frequencies of major leukocyte subsets across MB, mPLAC, fPLAC, and CB. **B** CD206$^{hi}$ CD14$^{+}$ monocyte/macrophage frequencies across MB, IVB, mPLAC or MB, decidua, and mPLAC. **C** Phenotypic differences between CD206$^{lo}$ and CD206$^{hi}$ CD14$^{+}$ macrophages/monocytes from mPLAC tissues. **A** depicts *n* = 26 dyads. **B** depicts *n* = 9 dyads. **C** depicts *n* = 22 dyads. Indicated are statistical significances where $p < 0.1$ by **A**, **B** Friedman tests with Dunn's multiple comparisons tests and **C** Wilcoxon tests. Symbols in **A**–**C** are connected by donor identity.

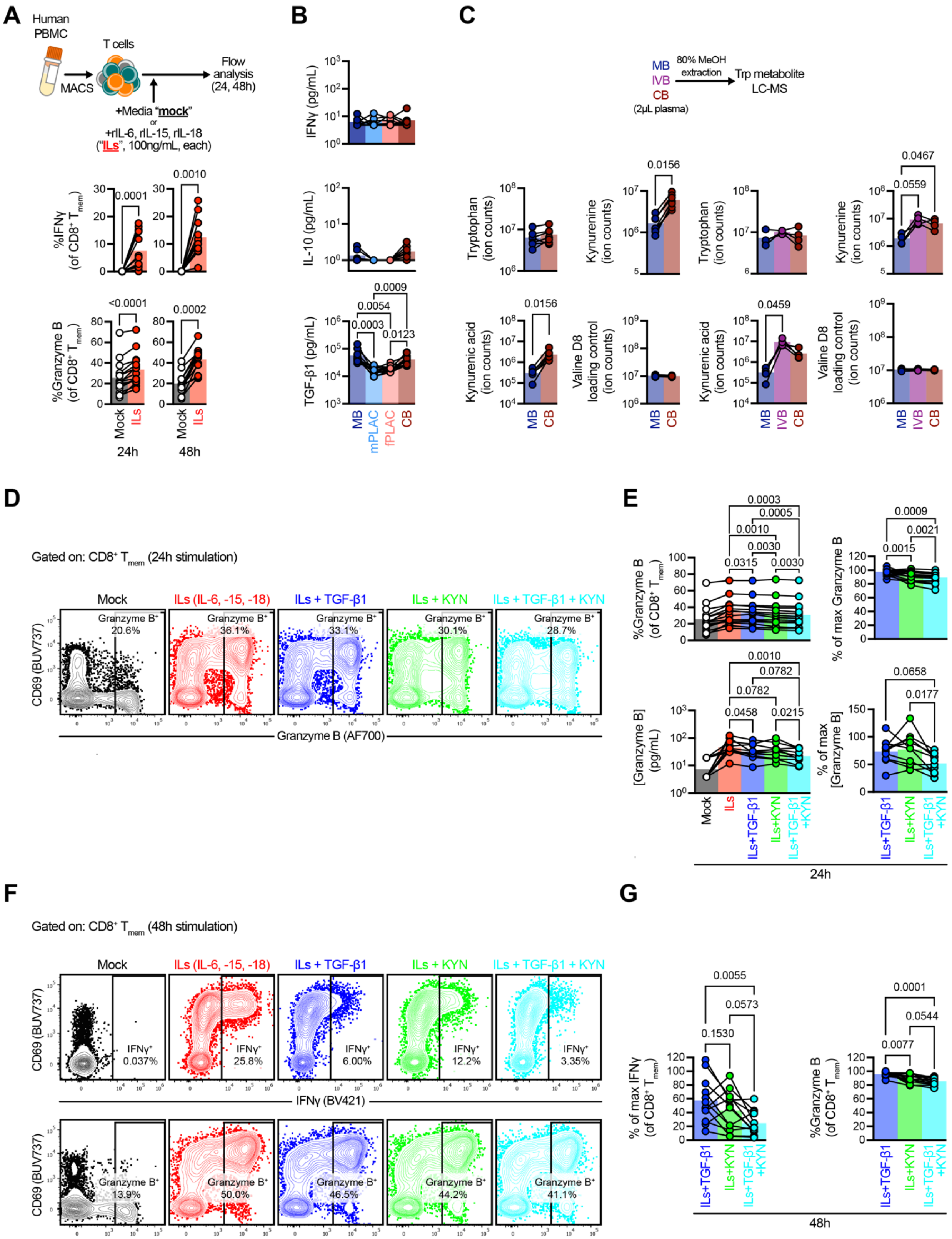
A
Human PBMC
MACS
T cells
Flow analysis (24, 48h)
+Media "mock"
or
+rIL-6, rIL-15, rIL-18 ("ILs", 100ng/mL, each)
%IFNγ (of CD8+ Tmem)
%Granzyme B (of CD8+ Tmem)
0.0001
0.0010
<0.0001
0.0002
Mock
ILs
24h
48h
B
IFNγ (pg/mL)
IL-10 (pg/mL)
TGF-β1 (pg/mL)
0.0009
0.0054
0.0003
0.0123
MB
mPLAC
fPLAC
CB
C
MB
IVB
CB
(2μL plasma)
80% MeOH extraction
Trp metabolite LC-MS
Tryptophan (ion counts)
Kynurenine (ion counts)
Kynurenic acid (ion counts)
Valine D8 loading control (ion counts)
0.0156
0.0467
0.0559
0.0459
D
Gated on: CD8+ Tmem (24h stimulation)
Mock
ILs (IL-6, -15, -18)
ILs + TGF-β1
ILs + KYN
ILs + TGF-β1 + KYN
Granzyme B+
20.6%
36.1%
33.1%
30.1%
28.7%
CD69 (BUV737)
Granzyme B (AF700)
E
%Granzyme B (of CD8+ Tmem)
% of max Granzyme B
[Granzyme B] (pg/mL)
% of max [Granzyme B]
0.0003
0.0005
0.0010
0.0030
0.0315
0.0030
0.0009
0.0021
0.0015
0.0010
0.0782
0.0782
0.0458
0.0215
0.0658
0.0177
Mock
ILs
ILs+TGF-β1
ILs+KYN
ILs+TGF-β1 +KYN
24h
F
Gated on: CD8+ Tmem (48h stimulation)
Mock
ILs (IL-6, -15, -18)
ILs + TGF-β1
ILs + KYN
ILs + TGF-β1 + KYN
IFNγ+
0.037%
25.8%
6.00%
12.2%
3.35%
IFNγ (BV421)
Granzyme B+
13.9%
50.0%
46.5%
44.2%
41.1%
CD69 (BUV737)
Granzyme B (AF700)
G
% of max IFNγ (of CD8+ Tmem)
%Granzyme B (of CD8+ Tmem)
0.0055
0.0573
0.1530
0.0001
0.0544
0.0077
ILs+TGF-β1
ILs+KYN
ILs+TGF-β1 +KYN
48h

**Supplemental figure 5**
**TGF-β1 and kynurenine impair inflammation-mediated IFNγ and GzmB upregulation.**

**A** Expression of IFNγ and GzmB after in vitro cytokine stimulation with IL-6, IL-15, and IL-18 (100ng/mL, each). **B** Concentrations of IFNγ, IL-10, and TGF-β1 across MB and CB plasmas and mPLAC and fPLAC lysates. **C** LC-MS for tryptophan metabolites and loading control in MB, IVB, and CB plasma. **D** Representative flow plots of GzmB expression in $CD8^+$ $T_{mem}$ across stimulation conditions after 24h. **E** Expression of GzmB in $CD8^+$ $T_{mem}$ (top) and concentration of GzmB in culture supernatant (bottom) across stimulation conditions after 24h. **F** Representative flow plots of IFNγ and GzmB expression in $CD8^+$ $T_{mem}$ across stimulation conditions after 48h. **G** Percent of maximal IFNγ (left) and GzmB (right) expression in $CD8^+$ $T_{mem}$ after 48h. **A** depicts $n$ = 15 PBMC donors. **B** Depicts $n$ = 8–12 dyads. **C** depicts $n$ = 4 dyads with IVB and $n$ = 7 MB and CB dyads. **D** depicts $n$ = 15 and $n$ = 10 PBMC donors analyzed via flow and multiplex cytokine analysis, respectively. **G** depicts $n$ = 12 PBMC donors. Symbols are connected by donor identity. Indicated are statistical significances where $p < 0.1$ by **A**–**C** Wilcoxon tests and Friedman tests with Dunn's multiple comparisons or **D**, **G** RM one-way ANOVA with Geisser-Greenhouse correction and Holm-Šídák multiple comparisons tests.

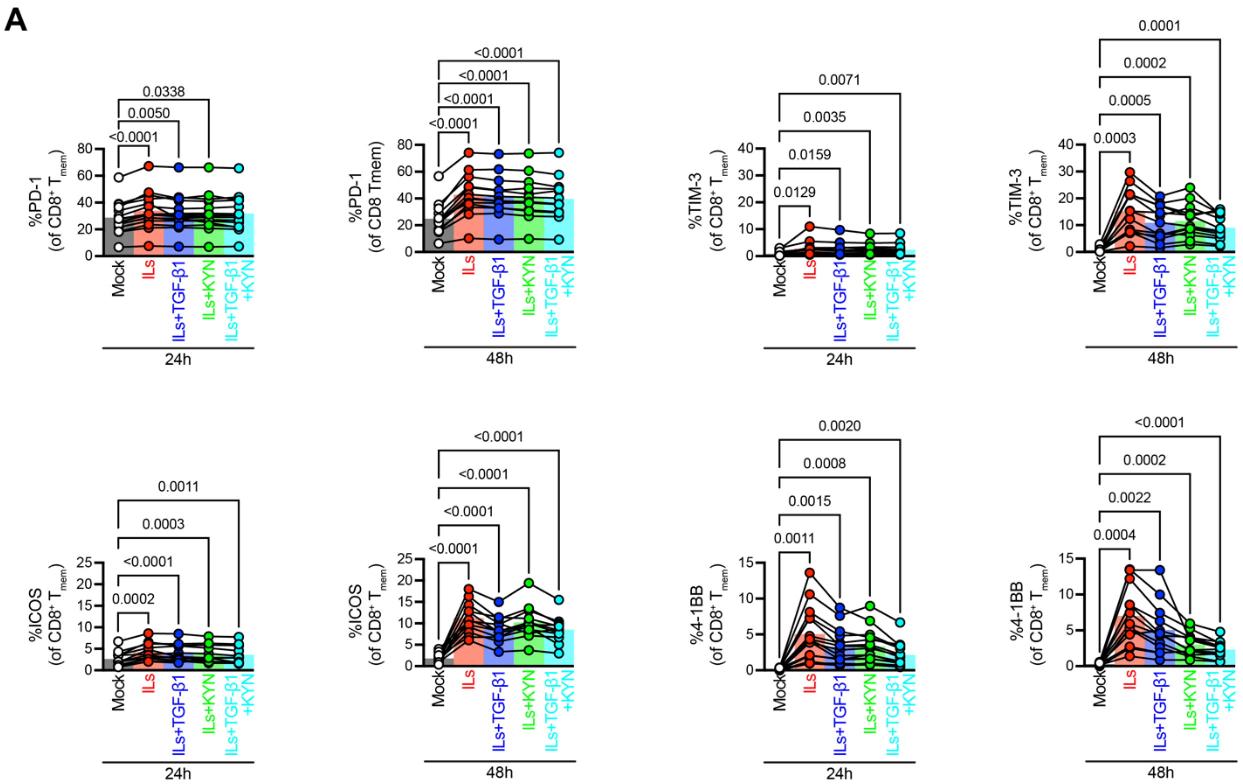


**Supplemental figure 6**

**TGF-β1 and kynurenine do not abrogate inflammation-mediated receptor upregulation in CD8+ $T_{mem}$**

**A** PD-1, TIM-3, ICOS, and 4-1BB expression frequencies in CD8+ $T_{mem}$ across stimulation conditions at 24h and 48h timepoints. **A** depicts *n* = 12–15 PBMC donors with symbols connected by donor identity. Indicated are statistical significance where $p < 0.1$ by RM one-way ANOVA with Geisser-Greenhouse correction and Holm-Šídák multiple comparisons tests (all columns compared to mock).

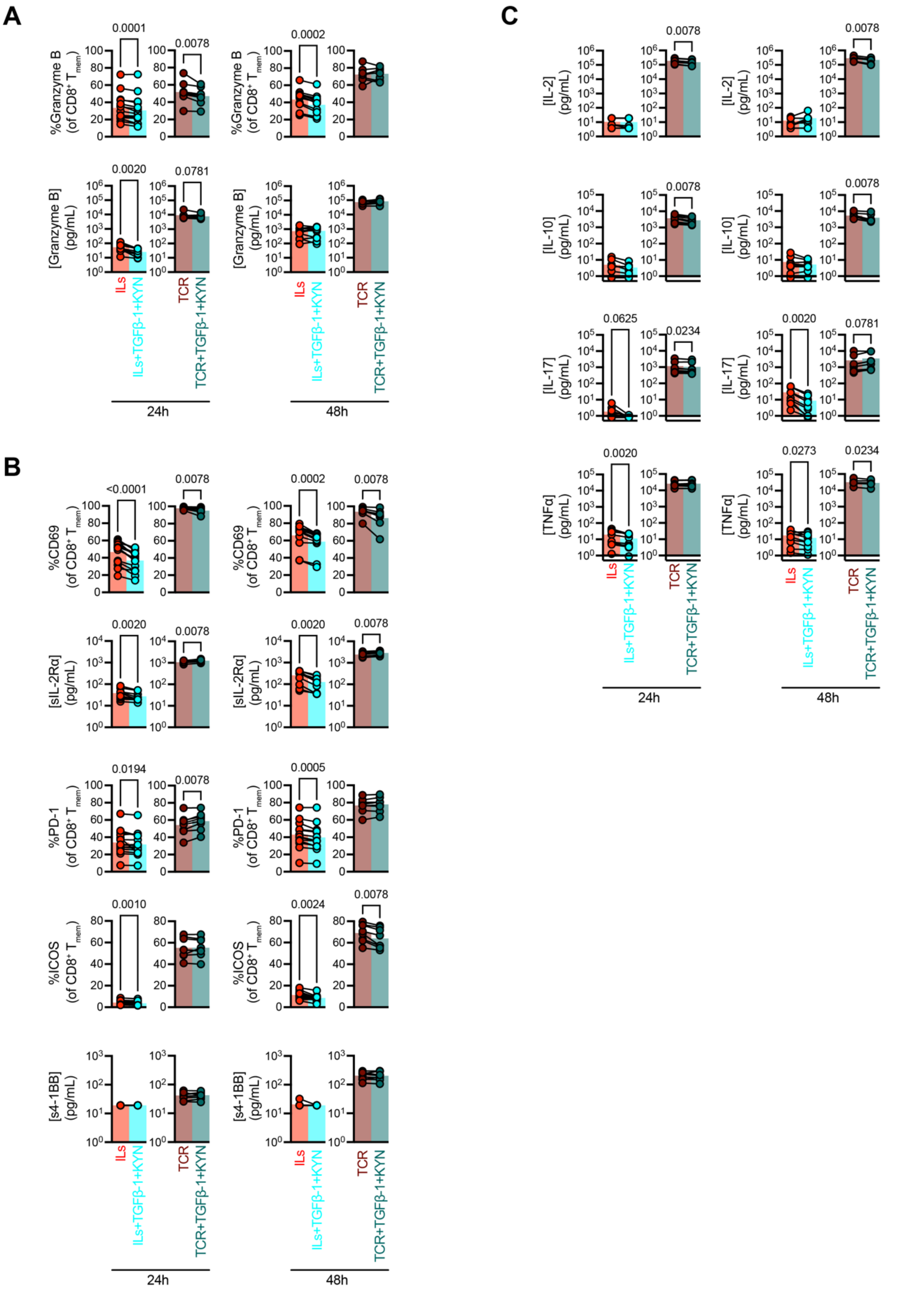


**Supplemental figure 7**
**Differential effects of TGF-β1 and kynurenine on TCR- and cytokine-mediated activation.**

Cells were stimulated as described in Figure 6B where MACS-isolated T cells were stimulated with IL-6, IL-15, IL-18 stimulation (“ILs”; 100ng/mL, each) or anti-CD3/CD28 microbead (“TCR”; 1:1 bead:cell ratio) in the presence or absence of TGF-β1 (0.2ng/mL) and kynurenine (100μM) for 24 and 48h. Afterwards, T cells were characterized via flow cytometry and culture supernatants were analyzed using multiplex cytokine analysis. **A** GzmB expression in $CD8^+$ $T_{mem}$ and concentration in culture supernatants. **B** Expression of activations markers (CD69, PD-1, ICOS) in $CD8^+$ $T_{mem}$ or concentrations of factors indicative of activation (sIL-2Rα, s4-1BB) across stimulation conditions. **C** Expression of effector molecules (IL-2, IL-10, IL-17, TNFα) in culture supernatants across stimulation conditions. **A**–**C** depict 8–15 PBMC donors, with symbols connected by donor identity. Indicated are statistical significance where $p < 0.1$ by Wilcoxon tests.

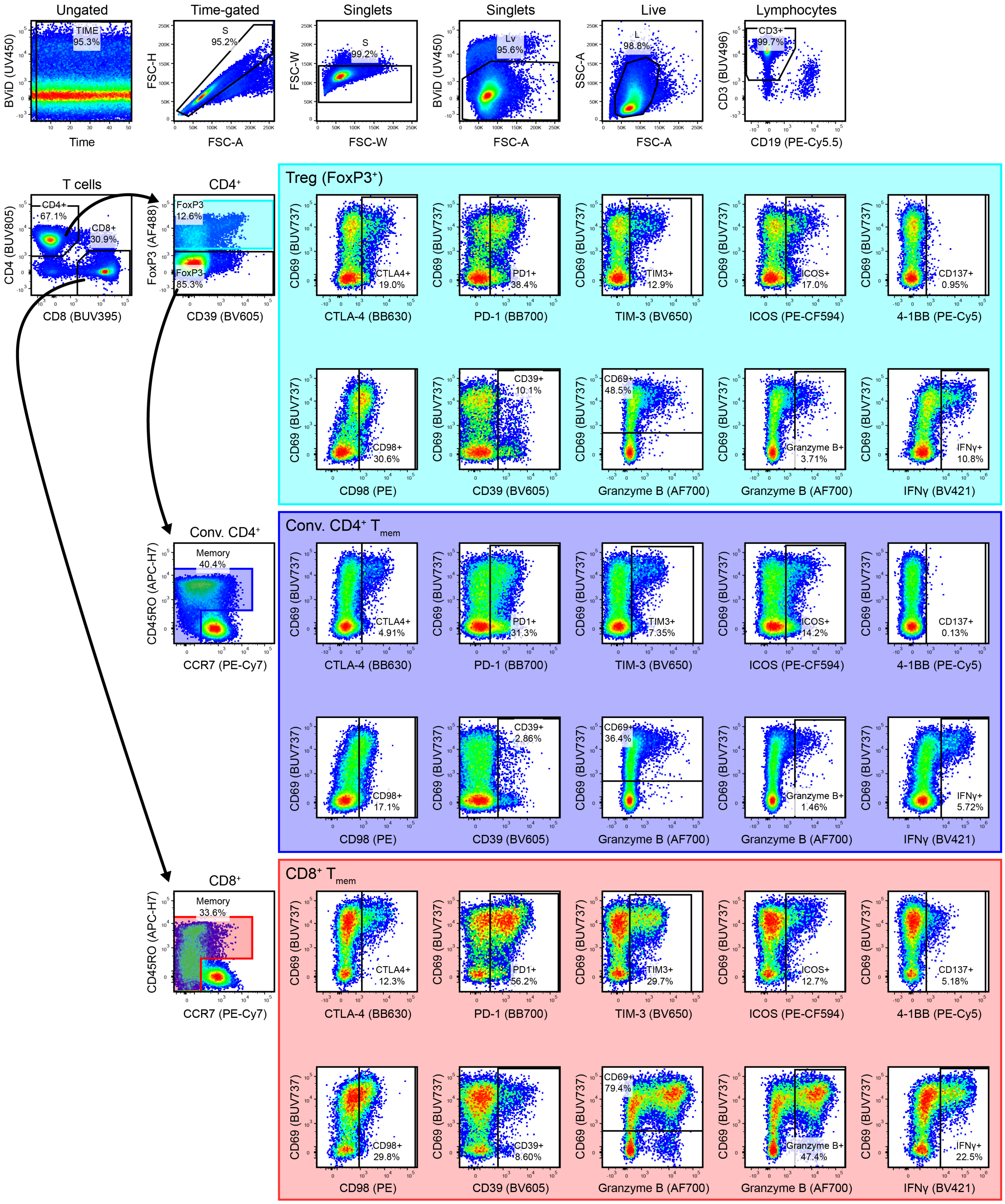


**Supplemental figure 8**
**Gating strategy for T cell stimulation and intracellular cytokine staining panel**

Gating strategy depicts T cells after stimulation with IL-6, IL-15, and IL-18 in combination (100ng/mL, each) for 48 h.

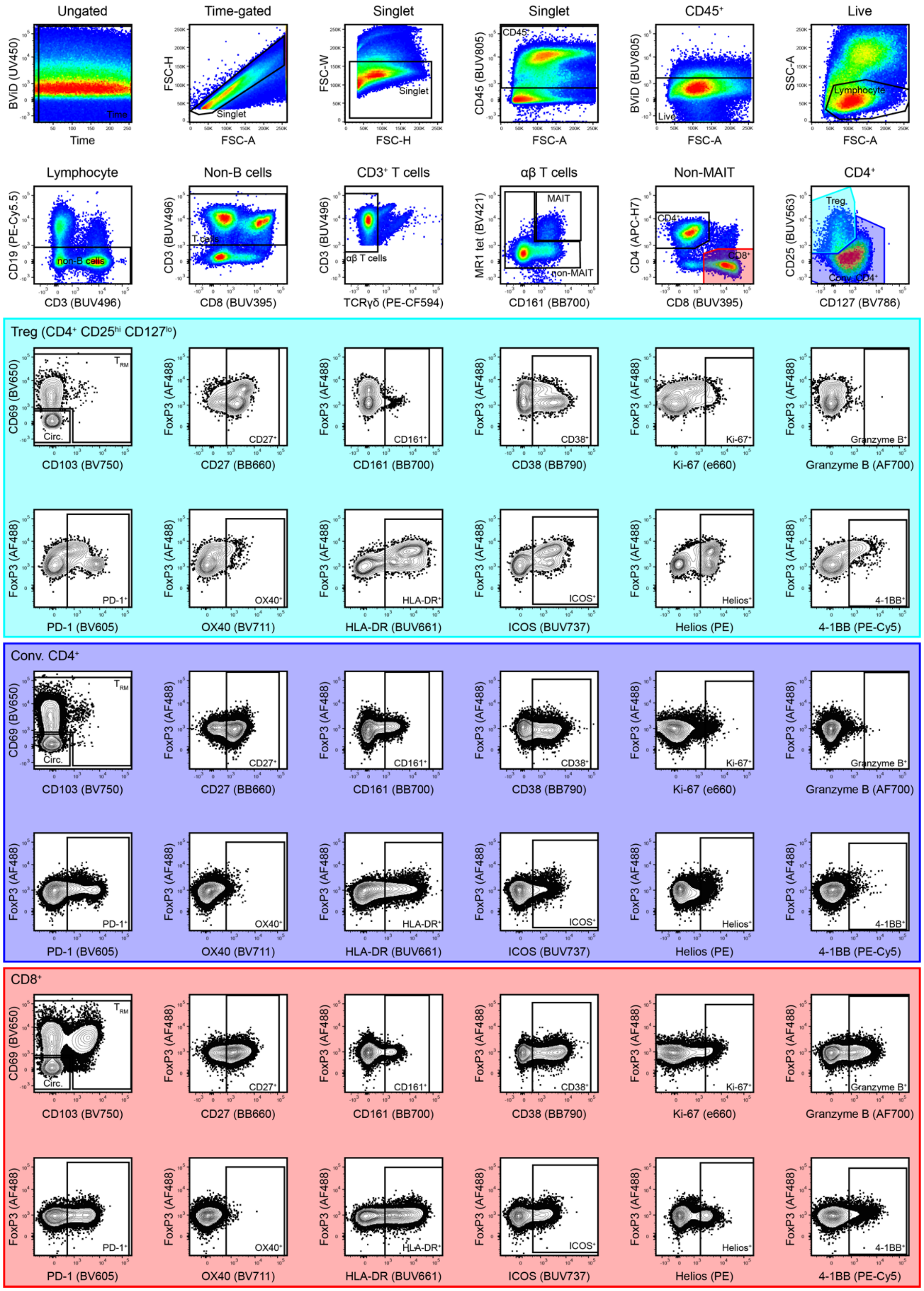


**Supplemental figure 9**
**Gating strategy for T cell phenotyping panel 1**

Representative gating example from cells freshly isolated from mPLAC

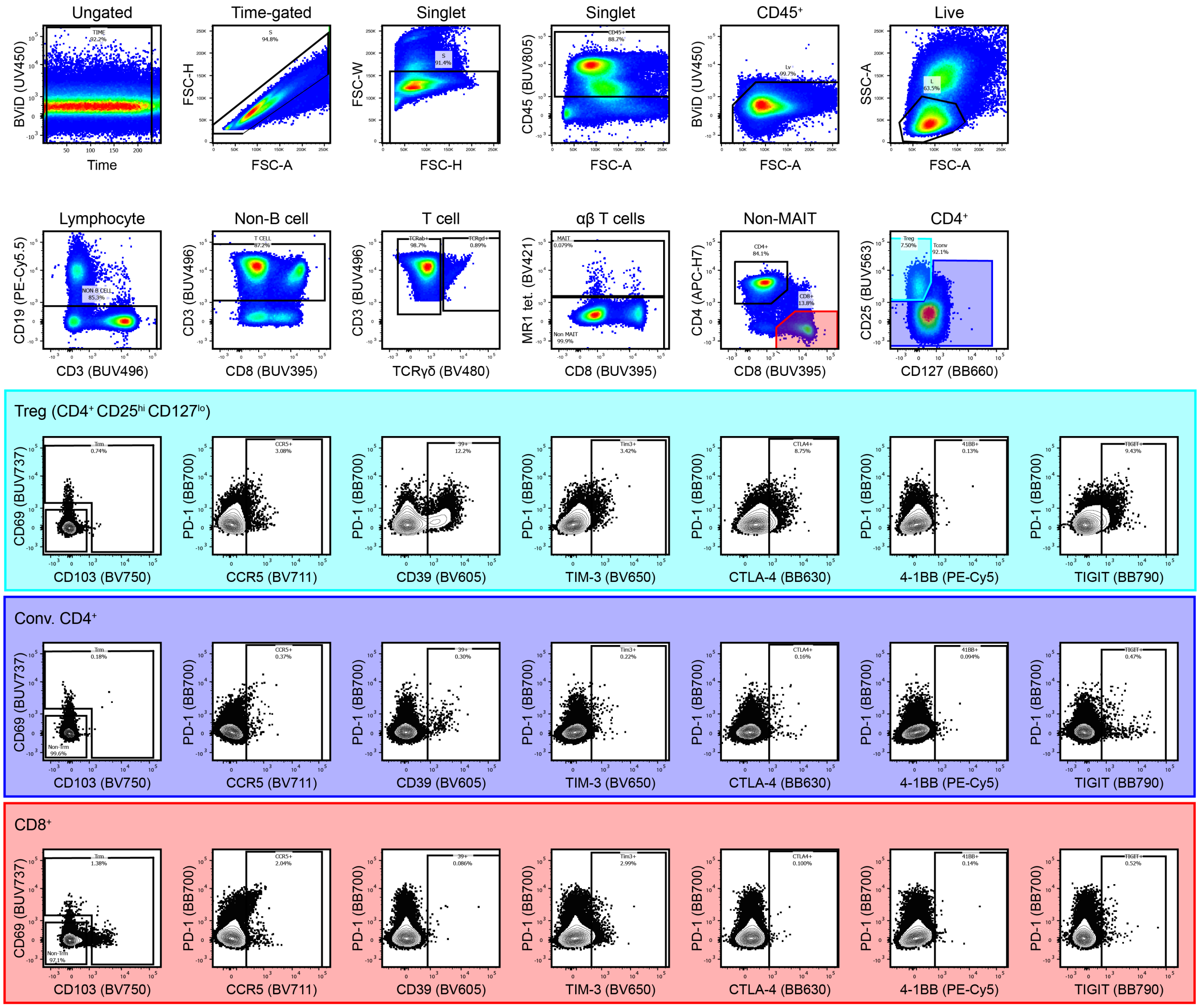


**Supplemental figure 10**
**Gating strategy for T cell phenotyping panel 2 (CB)**

Representative gating example from cell freshly isolated from CB

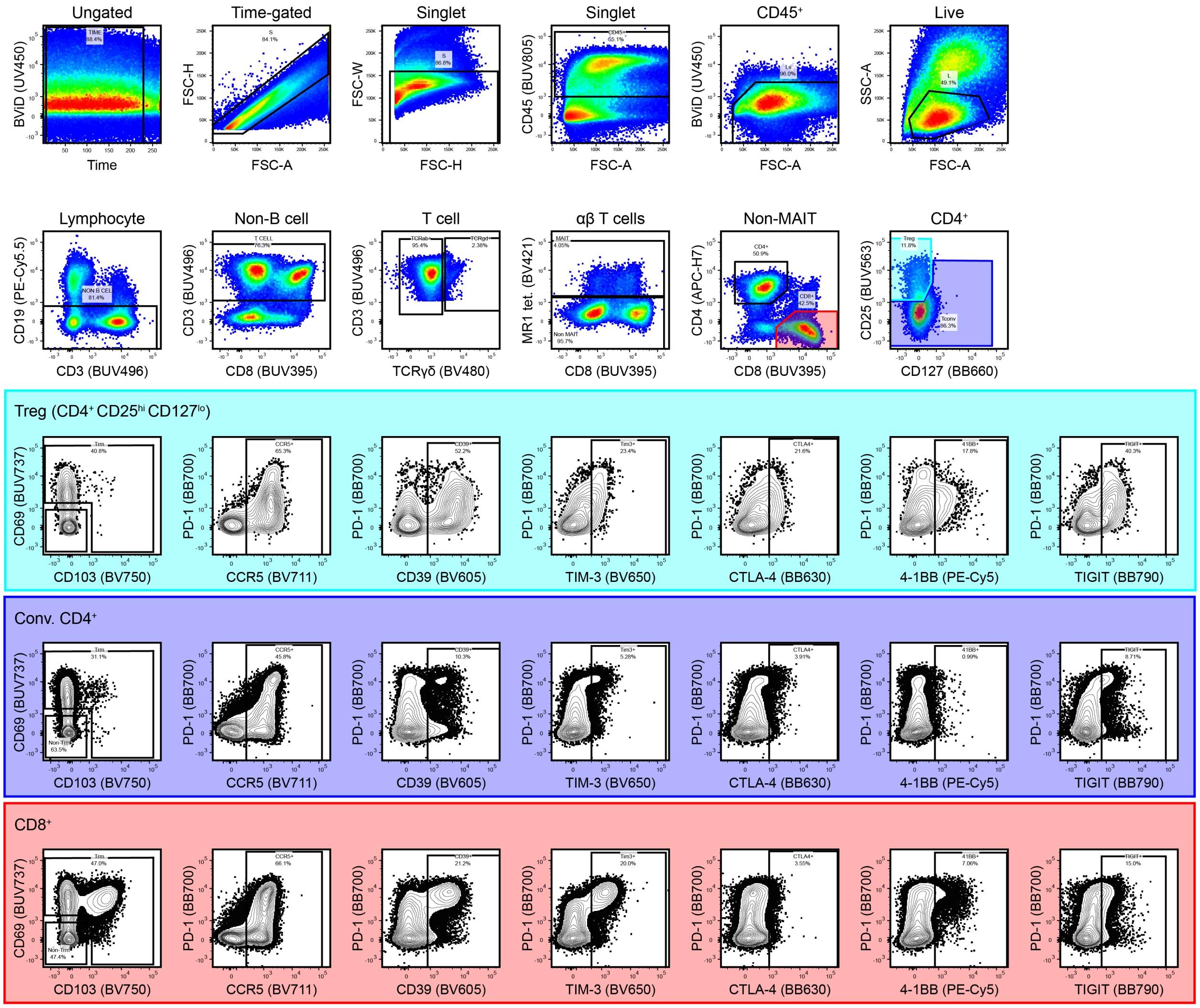


**Supplemental figure 11**
**Gating strategy for T cell phenotyping panel 2 (mPLAC)**

Representative gating example from cell freshly isolated from mPLAC

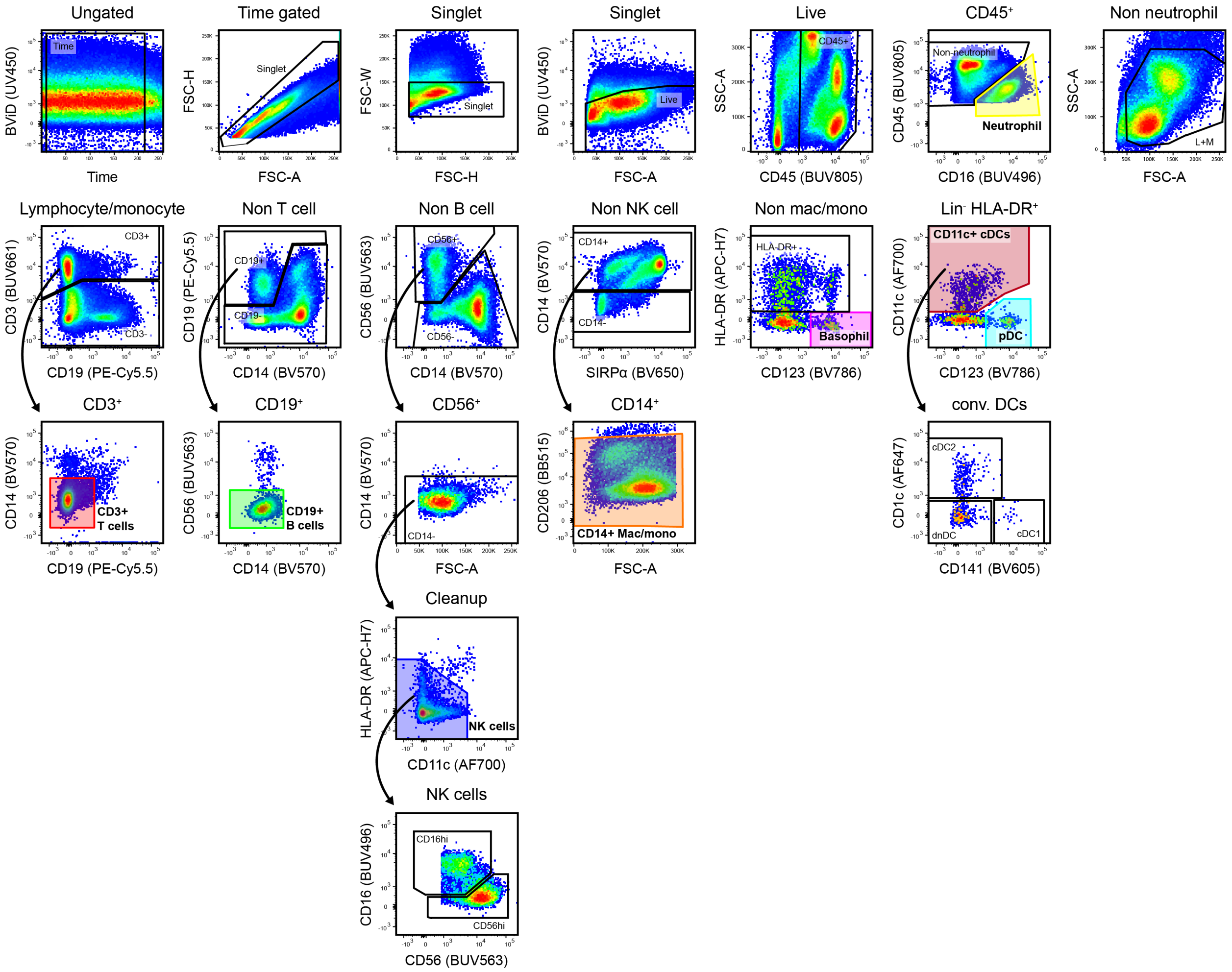


**Supplemental figure 12**
**Gating strategy for major leukocyte populations for APC panels 1 and 2**

Representative gating example for APC panels (shared backbone for major lineages) from cells freshly isolated from mPLAC

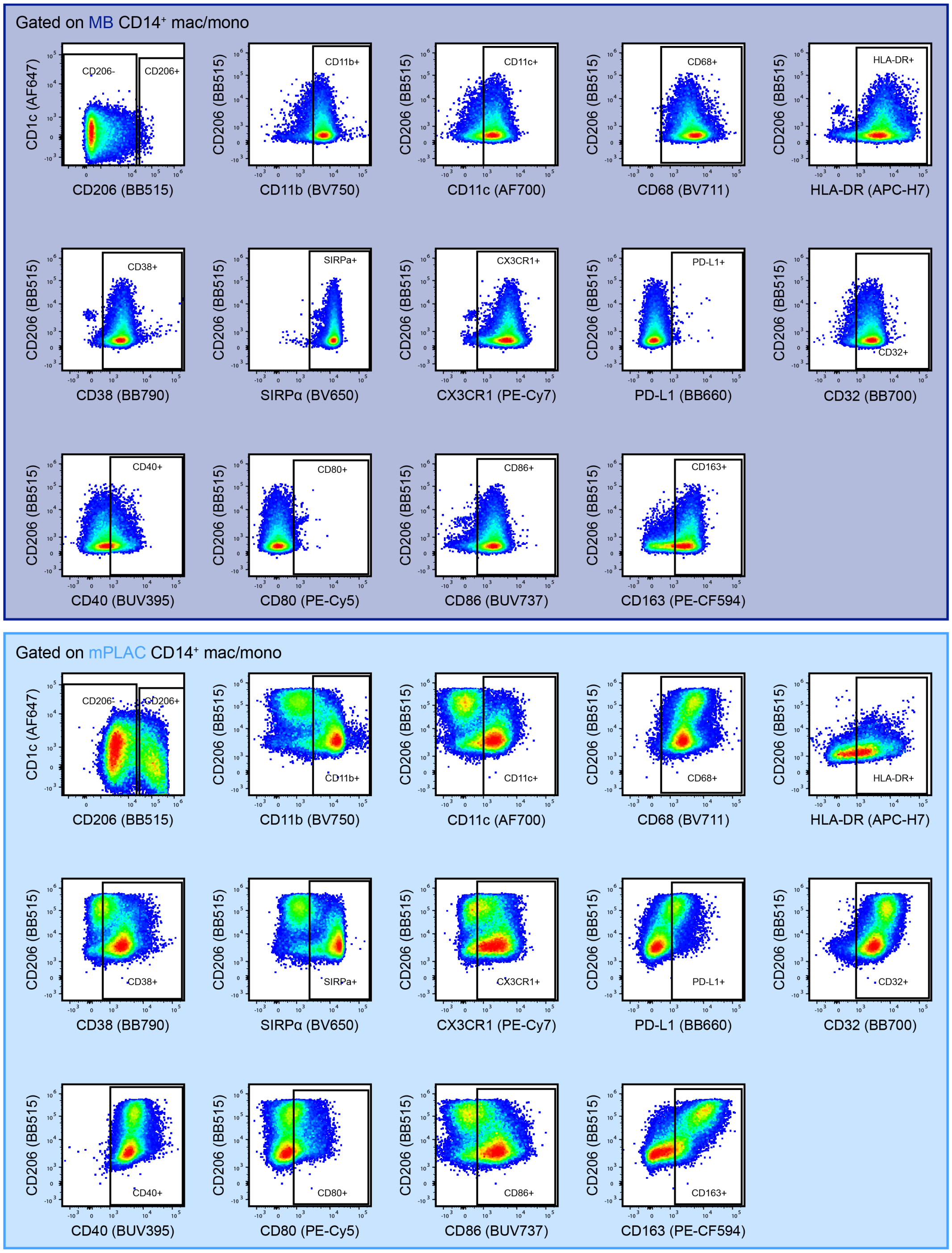


**Supplemental figure 13**
**APC panel 1 gating strategy for CD14$^+$ macrophage/monocyte phenotyping**

Representative gating example for CD14$^+$ macrophage/monocyte events from cells freshly isolated from MB (top) or mPLAC (bottom)

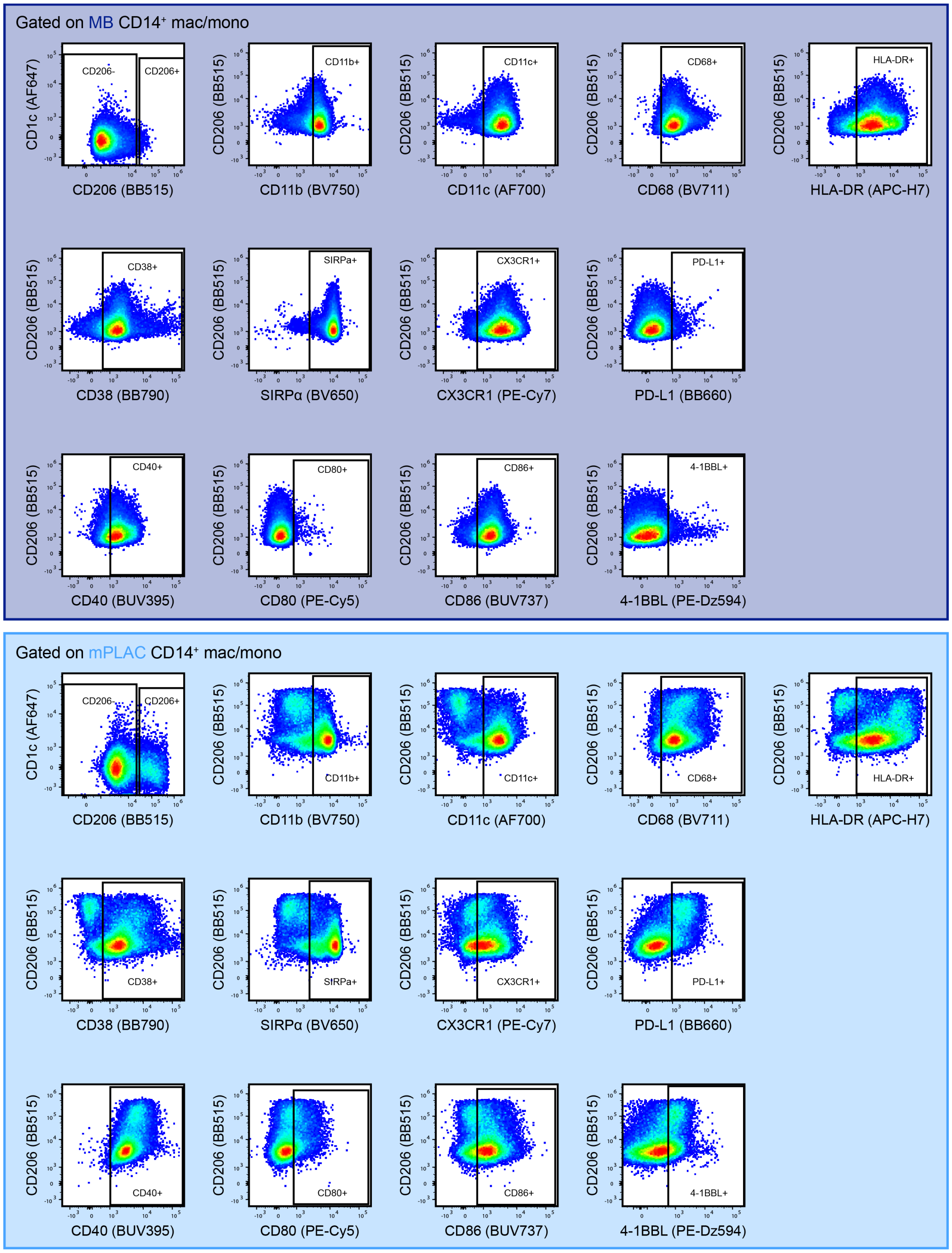


**Supplemental figure 14**
**APC panel 2 gating strategy for CD14$^+$ macrophage/monocyte phenotyping**

Representative gating example for CD14$^+$ macrophage/monocyte events from cells freshly isolated from MB (top) or mPLAC (bottom)

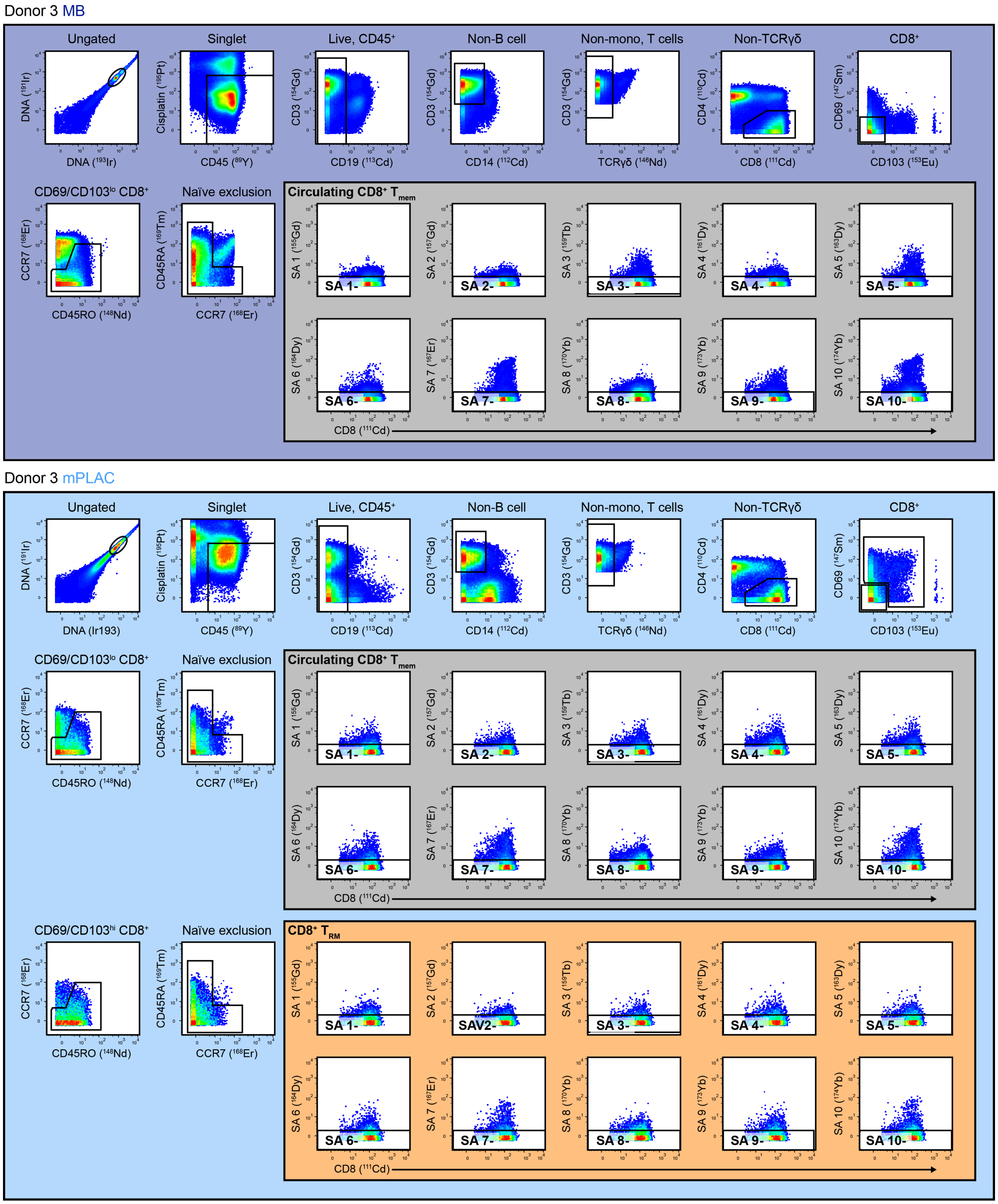


#### Ag-specific population identificaiton

Boolean gating combining all subset-specific events negative for all but two of surveyed SAs

**Gated on:**
SA 1-, SA 2-, SA 3-, SA 4-, SA 5-, SA 6-, SA 8-, SA 9-

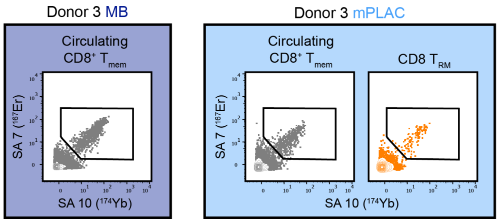


**Gated on:**
SA 1-, SA 2-, SA 3-, SA 4-, SA 5-, SA 6-, SA 8-, SA 10-

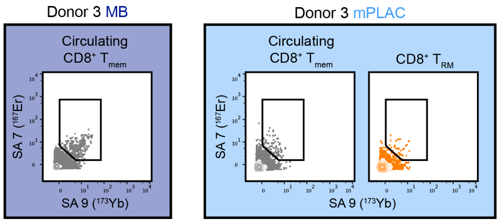

**Supplemental figure 15**
**Gating strategy for CyTOF tetramer screen staining panel**

We identified circulating memory (grey) and resident memory (orange) $CD8^+$ T cells in cryopreserved MB (dark blue) and mPLAC (light blue) samples. Within these subpopulations, we gated tetramer-negative populations, which we combined all but two (in all possible combinations) using Boolean gating. Within these Booleans, we measured tetramer-SA staining for the two excluded SAs to identify Ag-specific $CD8^+$ T cells within each tissue and memory subset.

**Supplemental table 1**
**Cell counting panel for Guava EasyCyte**

| Reagent | Conjugate | Clone | Vendor | Dilution |
|---|---|---|---|---|
| Guava stain (FACSWash diluent, 20 min, room temperature) | | | | |
| 7-AAD | NA | NA | BD Biosciences | 1:15 |
| CD45 | FITC | 2D1 | BD Biosciences | 1:50 |
| CD3 | PE | UCHT1 | BD Biosciences | 1:30 |
| Fix (1% PFA) | | | | |

**Supplemental table 2**
**T cell stimulation and intracellular cytokine staining panel**

| Reagent | Conjugate | Clone | Vendor | Dilution |
|---|---|---|---|---|
| Viability stain (1x PBS diluent, 20 min, room temperature) | | | | |
| LIVE/DEAD fixable blue viability dye (BViD) | UV450 | NA | Thermo Fisher | 1:500 |
| Surface stain (FACSwash + Azide diluent, 20 min, room temperature) | | | | |
| Brilliant stain buffer plus | NA | NA | BD Biosciences | 1:5 |
| TruStain FcX (Fc block) | NA | NA | BioLegend | 1:20 |
| CD69 | BUV737 | FN50 | BD Biosciences | 1:40 |
| CD39 | BV605 | A1 | BioLegend | 1:20 |
| TIM-3 | BV650 | 7D3 | BD Biosciences | 1:20 |
| PD-1 | BB700 | EH12.1 | BD Biosciences | 1:20 |
| CD98 | PE | MEM-108 | Thermo Fisher | 1:100 |
| CD278 (ICOS) | PE-CF594 | C398.4A | BD Biosciences | 1:1000 |
| CD137 (4-1BB) | PE-Cy5 | 4B4-1 | BD Biosciences | 1:20 |
| CD19 | PE-Cy5.5 | SJ25-C1 | Thermo Fisher | 1:160 |
| CCR7 | PE-Cy7 | 3D12 | BD Biosciences | 1:40 |
| CD45RO | APC-H7 | UCHL1 | BD Biosciences | 1:40 |
| Fix (1x eBioscience FOXP3 fixation buffer, 20 min, room temperature) | | | | |
| Intracellular stain (1x eBioscience FOXP3 permeabilization buffer, 30 min, room temperature) | | | | |
| CD8 | BUV395 | RPA-T8 | BD Biosciences | 1:80 |
| CD3 | BUV496 | UCHT1 | BD Biosciences | 1:40 |
| CD4 | BUV805 | SK3 | BD Biosciences | 1:80 |
| IFNγ | BV421 | 4S.B3 | BioLegend | 1:40 |
| FoxP3 | AF488 | 259D/C7 | BD Biosciences | 1:10 |
| CTLA-4 | BB630 | BNI3 | BD Biosciences | 1:80 |
| TOX | APC | REA473 | Miltenyi Biotec | 1:80 |
| Granzyme B | AF700 | GB11 | BD Biosciences | 1:80 |

**Supplemental table 3**
**FACS panel for scRNAseq**

| Reagent | Conjugate | Clone | Vendor | Dilution |
|---|---|---|---|---|
| Viability stain (1x PBS diluent, 20 min, room temperature) | | | | |
| LIVE/DEAD fixable aqua viability dye (AViD) | V510 | NA | Thermo Fisher | 1:500 |
| Tetramer stain (FACSwash diluent, 45 min, room temperature) | | | | |
| TruStain FcX (Fc block) | NA | NA | BioLegend | 1:20 |
| MR1 OP-5-RU (MAIT cell) tetramer | PE | NA | NIH Tetramer core | 1:400 |
| Surface stain (FACSwash diluent, 20 min, room temperature) | | | | |
| CD25 | BV421 | M-A251 | BD Biosciences | 1:40 |
| CD14 | BV711 | MφP9 | BD Biosciences | 1:40 |
| CD127 | BV786 | HIL-7R-M21 | BD Biosciences | 1:10 |
| CD45 | FITC | 2D1 | BD Biosciences | 1:40 |
| CD19 | PE-Cy5.5 | SJ25-C1 | Thermo Fisher | 1:160 |
| CD8 | PE-Cy7 | RPA-T8 | BD Biosciences | 1:40 |
| CD3 | APC | SK7 | BD Biosciences | 1:50 |
| CD4 | AF700 | RPA-T4 | BD Biosciences | 1:100 |
| HLA-DR | APC-H7 | G46-6 | BD Biosciences | 1:40 |

**Supplemental table 4**
**T cell phenotyping panel 1**

| Reagent | Conjugate | Clone | Vendor | Dilution |
|---|---|---|---|---|
| Viability stain (1x PBS diluent, 20 min, room temperature) | | | | |
| LIVE/DEAD fixable blue viability dye (BViD) | UV450 | NA | Thermo Fisher | 1:500 |
| Tetramer stain (FACSWash + Azide diluent, 60 min, room temperature) | | | | |
| MR1 OP-5-RU (MAIT cell) tetramer | BV421 | NA | NIH Tetramer Core | 1:500 |
| TruStain FcX (Fc block) | NA | NA | BioLegend | 1:20 |
| Surface stain (FACSwash + Azide diluent, 20 min, room temperature) | | | | |
| Brilliant stain buffer concentrate | NA | NA | BD Biosciences | 1:5 |
| CD8 | BUV395 | RPA-T8 | BD Biosciences | 1:80 |
| CD3 | BUV496 | UCHT1 | BD Biosciences | 1:40 |
| CD25 | BUV563 | 2A3 | BD Biosciences | 1:40 |
| HLA-DR | BUV661 | G46-6 | BD Biosciences | 1:80 |
| CD278 (ICOS) | BUV737 | DX29 | BD Biosciences | 1:10 |
| CD45 | BUV805 | HI30 | BD Biosciences | 1:80 |
| CD28 | BV480 | CD28.2 | BD Biosciences | 1:40 |
| CD45RA | BV570 | HI100 | BioLegend | 1:160 |
| PD-1 | BV605 | EH12.1 | BD Biosciences | 1:20 |
| CD69 | BV650 | FN50 | BD Biosciences | 1:20-40* |
| OX40 | BV711 | Act35 | BD Biosciences | 1:40 |
| CD103 | BV750 | Ber-ACT8 | BD Biosciences | 1:160 |
| CD127 | BV786 | HIL-7R-M21 | BD Biosciences | 1:10 |
| CD27 | BB660 | M-T271 | BD Biosciences | 1:160 |
| CD161 | BB700 | DX12 | BD Biosciences | 1:20 |
| CD38 | BB790 | HIT2 | BD Biosciences | 1:80 |
| TCRγδ | PE-CF594 | B1 | BD Biosciences | 1:20 |
| CD137 (4-1BB) | PE-Cy5 | 4B4-1 | BD Biosciences | 1:20 |
| CD19 | PE-Cy5.5 | SJ25-C1 | Thermo Fisher | 1:160 |
| CCR7 | PE-Cy7 | 3D12 | BD Biosciences | 1:40 |
| CD4 | APC-H7 | RPA-T4 | BD Biosciences | 1:40 |
| Fix (1x eBioscience FOXP3 fixation buffer, 20 min, room temperature) | | | | |
| Intracellular stain (1x eBioscience FOXP3 permeabilization buffer, 30 min, room temperature) | | | | |
| FoxP3 | AF488 | 259D/C7 | BD Biosciences | 1:20 |
| CTLA-4 | BB630 | BNI3 | BD Biosciences | 1:80 |
| Helios | PE | 22F6 | BD Biosciences | 1:500 |
| Ki-67 | e660 | Sola15 | Thermo Fisher | 1:1000 |
| Granzyme B | AF700 | GB11 | BD Biosciences | 1:80 |

*Lot-dependent dilution variability

**Supplemental table 5**
**T cell phenotyping panel 2**

| Reagent | Conjugate | Clone | Vendor | Dilution |
|---|---|---|---|---|
| Viability stain (1x PBS diluent, 20 min, room temperature) | | | | |
| LIVE/DEAD fixable blue viability dye (BViD) | UV450 | NA | Thermo Fisher | 1:500 |
| Tetramer stain (FACSWash + Azide diluent, 60 min, room temperature) | | | | |
| MR1 OP-5-RU (MAIT cell) tetramer | BV421 | NA | NIH Tetramer Core | 1:500 |
| TruStain FcX (Fc block) | NA | NA | BioLegend | 1:20 |
| 1º Surface stain (FACSwash + Azide diluent, 20 min, room temperature) | | | | |
| CD127 | Biotin | A019D5 | BioLegend | 1:40 |
| 2º Surface stain (FACSwash + Azide diluent, 20 min, room temperature) | | | | |
| Brilliant stain buffer plus | NA | NA | BD Biosciences | 1:5 |
| CD8 | BUV395 | RPA-T8 | BD Biosciences | 1:80 |
| CD3 | BUV496 | UCHT1 | BD Biosciences | 1:40 |
| CD25 | BUV563 | 2A3 | BD Biosciences | 1:40 |
| CD69 | BUV737 | FN50 | BD Biosciences | 1:80 |
| CD45 | BUV805 | HI30 | BD Biosciences | 1:80 |
| TCRγδ | BV480 | B1 | BD Biosciences | 1:10 |
| CD45RA | BV570 | HI100 | BioLegend | 1:160 |
| CD39 | BV605 | A1 | BioLegend | 1:20 |
| TIM-3 | BV650 | 7D3 | BD Biosciences | 1:20 |
| CCR5 | BV711 | 2D7/CCR5 | BD Biosciences | 1:20 |
| CD103 | BV750 | Ber-ACT8 | BD Biosciences | 1:160 |
| Ki-67 | BV786 | B56 | BD Biosciences | 1:320 |
| Streptavidin | BB660 | NA | BD Biosciences | 1:400 |
| PD-1 | BB700 | EH12.1 | BD Biosciences | 1:20 |
| TIGIT | BB790 | 741182 | BD Biosciences | 1:80 |
| CD137 (4-1BB) | PE-Cy5 | 4B4-1 | BD Biosciences | 1:20 |
| CD19 | PE-Cy5.5 | SJ25-C1 | Thermo Fisher | 1:160 |
| CD4 | APC-H7 | RPA-T4 | BD Biosciences | 1:40 |
| Fix (1x eBioscience FOXP3 fixation buffer, 20 min, room temperature) | | | | |
| Intracellular stain (1x eBioscience FOXP3 permeabilization buffer, 30 min, room temperature) | | | | |
| FoxP3 | AF488 | 259D/C7 | BD Biosciences | 1:20 |
| CTLA-4 | BB630 | BNI3 | BD Biosciences | 1:80 |
| TCF-1 | PE | 22F6 | BD Biosciences | 1:500 |
| Eomes | PE-e610 | WD1928 | Thermo Fisher | 1:20 |
| Tbet | PE-Cy7 | 4B10 | Thermo Fisher | 1:40 |
| TOX | APC | REA473 | Miltenyi Biotec | 1:80 |
| Granzyme B | AF700 | GB11 | BD Biosciences | 1:80 |

**Supplemental table 6**
**T cell phenotyping panel 3**

| Reagent | Conjugate | Clone | Vendor | Dilution |
|---|---|---|---|---|
| Viability stain (1x PBS diluent, 20 min, room temperature) | | | | |
| LIVE/DEAD fixable blue viability dye (BViD) | UV450 | NA | Thermo Fisher | 1:500 |
| 1º Surface stain (FACSwash + Azide diluent, 20 min, room temperature) | | | | |
| CD127 | Biotin | A019D5 | BioLegend | 1:40 |
| TruStain FcX (Fc block) | NA | NA | BioLegend | 1:20 |
| 2º Surface stain (FACSwash + Azide diluent, 20 min, room temperature) | | | | |
| Brilliant stain buffer plus | NA | NA | BD Biosciences | 1:5 |
| CD8 | BUV395 | RPA-T8 | BD Biosciences | 1:80 |
| CD3 | BUV496 | UCHT1 | BD Biosciences | 1:40 |
| CCR7 | BUV661 | 2-L1-A | BD Biosciences | 1:80 |
| ICOS | BUV737 | DX29 | BD Biosciences | 1:20 |
| CD45 | BUV805 | HI30 | BD Biosciences | 1:80 |
| CD25 | BV421 | M-A251 | BD Biosciences | 1:40 |
| CD28 | BV480 | CD28.2 | BD Biosciences | 1:40 |
| CD45RA | BV570 | HI100 | BioLegend | 1:160 |
| CD39 | BV605 | A1 | BioLegend | 1:20 |
| CD69 | BV650 | FN50 | BD Biosciences | 1:20-40* |
| OX40 | BV711 | ACT35 | BD Biosciences | 1:40 |
| CD103 | BV750 | Ber-ACT8 | BD Biosciences | 1:160 |
| CCR5 | BV786 | 3A9 | BD Biosciences | 1:20 |
| TIM-3 | BB515 | 7D3 | BD Biosciences | 1:80 |
| Streptavidin | BB660 | NA | BD Biosciences | 1:400 |
| PD-1 | BB700 | EH12.1 | BD Biosciences | 1:20 |
| TIGIT | BB790 | 741182 | BD Biosciences | 1:80 |
| IL-1R1 | PE | Goat polyclonal | R&D Systems | 1:20 |
| CXCR3 | PE-CF594 | 1C6/CXCR3 | BD Biosciences | 1:20 |
| CD137 (4-1BB) | PE-Cy5 | 4B4-1 | BD Biosciences | 1:20 |
| CD19 | PE-Cy5.5 | SJ25-C1 | Thermo Fisher | 1:160 |
| IL-18Rα | PE-Cy7 | H44 | BioLegend | 1:40 |
| IL-1R2 | APC | 34141 | R&D Systems | 1:20 |
| HLA-DR | APC-R700 | G46-6 | BD Biosciences | 1:160 |
| CD4 | APC-H7 | RPA-T4 | BD Biosciences | 1:40 |
| Fix (1x eBioscience FOXP3 fixation buffer, 20 min, room temperature) | | | | |
| Intracellular stain (1x eBioscience FOXP3 permeabilization buffer, 30 min, room temperature) | | | | |
| CTLA-4 | BB630 | BNI3 | BD Biosciences | 1:80 |

*Lot-dependent dilution variability

**Supplemental table 6**
**T cell phenotyping panel 4**

| Reagent | Conjugate | Clone | Vendor | Dilution |
|---|---|---|---|---|
| Viability stain (1x PBS diluent, 20 min, room temperature) | | | | |
| LIVE/DEAD fixable blue viability dye (BViD) | UV450 | NA | Thermo Fisher | 1:500 |
| 1º Surface stain (FACSwash + Azide diluent, 20 min, room temperature) | | | | |
| CD127 | Biotin | A019D5 | BioLegend | 1:40 |
| TruStain FcX (Fc block) | NA | NA | BioLegend | 1:20 |
| 2º Surface stain (FACSwash + Azide diluent, 20 min, room temperature) | | | | |
| Brilliant stain buffer plus | NA | NA | BD Biosciences | 1:5 |
| CD8 | BUV395 | RPA-T8 | BD Biosciences | 1:80 |
| CD3 | BUV496 | UCHT1 | BD Biosciences | 1:40 |
| CCR7 | BUV661 | 2-L1-A | BD Biosciences | 1:80 |
| ICOS | BUV737 | DX29 | BD Biosciences | 1:20 |
| CD45 | BUV805 | HI30 | BD Biosciences | 1:80 |
| CD25 | BV421 | M-A251 | BD Biosciences | 1:40 |
| CD28 | BV480 | CD28.2 | BD Biosciences | 1:40 |
| CD45RA | BV570 | HI100 | BioLegend | 1:160 |
| CD39 | BV605 | A1 | BioLegend | 1:20 |
| CD69 | BV650 | FN50 | BD Biosciences | 1:20-40* |
| OX40 | BV711 | ACT35 | BD Biosciences | 1:40 |
| CD103 | BV750 | Ber-ACT8 | BD Biosciences | 1:160 |
| CCR5 | BV786 | 3A9 | BD Biosciences | 1:20 |
| TIM-3 | BB515 | 7D3 | BD Biosciences | 1:80 |
| Streptavidin | BB660 | NA | BD Biosciences | 1:400 |
| PD-1 | BB700 | EH12.1 | BD Biosciences | 1:20 |
| CD98 | PE | MEM-108 | Thermo Fisher | 1:100 |
| CXCR3 | PE-CF594 | 1C6/CXCR3 | BD Biosciences | 1:20 |
| CD137 (4-1BB) | PE-Cy5 | 4B4-1 | BD Biosciences | 1:20 |
| CD19 | PE-Cy5.5 | SJ25-C1 | Thermo Fisher | 1:160 |
| IL-18Rα | PE-Cy7 | H44 | BioLegend | 1:40 |
| HLA-DR | APC-R700 | G46-6 | BD Biosciences | 1:160 |
| CD4 | APC-H7 | RPA-T4 | BD Biosciences | 1:40 |
| Fix (1x eBioscience FOXP3 fixation buffer, 20 min, room temperature) | | | | |
| Intracellular stain (1x eBioscience FOXP3 permeabilization buffer, 30 min, room temperature) | | | | |
| CTLA-4 | BB630 | BNI3 | BD Biosciences | 1:80 |
| AhR | e660 | 4MEJJ | Thermo Fisher | 1:100 |

*Lot-dependent dilution variability

**Supplemental table 7**
**APC phenotyping panel 1**

| Reagent | Conjugate | Clone | Vendor | Dilution |
|---|---|---|---|---|
| Viability stain (1x PBS diluent, 20 min, room temperature) | | | | |
| LIVE/DEAD fixable blue viability dye (BViD) | UV450 | NA | Thermo Fisher | 1:500 |
| 1º Surface stain (FACSwash + Azide diluent, 20 min, room temperature) | | | | |
| CX3CR1 | PE-Cy7 | 2A9-1 | BioLegend | 1:160 |
| TruStain FcX (Fc block) | NA | NA | BioLegend | 1:20 |
| 2º Surface stain (FACSwash + Azide diluent, 20 min, room temperature) | | | | |
| Brilliant stain buffer concentrate | NA | NA | BD Biosciences | 1:5 |
| CD40 | BUV395 | 5C3 | BD Biosciences | 1:40 |
| CD16 | BUV496 | 3G8 | BD Biosciences | 1:320 |
| CD56 | BUV563 | NCAM16.2 | BD Biosciences | 1:160 |
| CD3 | BUV661 | UCHT1 | BD Biosciences | 1:80 |
| CD86 | BUV737 | 2331 | BD Biosciences | 1:40 |
| CD45 | BUV805 | HI30 | BD Biosciences | 1:80 |
| PD-L2 | BV421 | MIH18 | BD Biosciences | 1:20 |
| CD85k | BV480 | ZM3.8 | BD Biosciences | 1:40 |
| CD14 | BV570 | M5E2 | BioLegend | 1:20 |
| CD141 | BV605 | 1A4 | BD Biosciences | 1:640 |
| SIRPα | BV650 | SE5A5 | BD Biosciences | 1:160 |
| CD11b | BV750 | ICRF44 | BD Biosciences | 1:160 |
| CD123 | BV786 | 7G3 | BD Biosciences | 1:40 |
| CD206 | BB515 | 19.2 | BD Biosciences | 1:20 |
| BTLA | BB630 | J168-540 | BD Biosciences | 1:80 |
| PD-L1 | BB660 | MIH1 | BD Biosciences | 1:40 |
| CD32 | BB700 | FLI8.26 | BD Biosciences | 1:160 |
| CD38 | BB790 | HIT2 | BD Biosciences | 1:80 |
| Axl | PE | 108724 | R&D Systems | 1:20 |
| CD163 | PE-CF594 | GHI/61 | BD Biosciences | 1:40 |
| CD80 | PE-Cy5 | L37.4 | BD Biosciences | 1:10 |
| CD19 | PE-Cy5.5 | SJ25-C1 | Thermo Fisher | 1:160 |
| CD1c | AF647 | F10/21A3 | BD Biosciences | 1:160 |
| CD11c | AF700 | B-ly6 | BD Biosciences | 1:320 |
| HLA-DR | APC-H7 | G46-6 | BD Biosciences | 1:40 |
| Fix (1x BD Cytofix/Cytoperm, 20 min, room temperature) | | | | |
| Intracellular stain (1x BD Perm/Wash buffer, 30 min, room temperature) | | | | |
| CD68 | BV711 | Y1/82A | BD Biosciences | 1:40 |

**Supplemental table 8**
**APC phenotyping panel 2**

| Reagent | Conjugate | Clone | Vendor | Dilution |
|---|---|---|---|---|
| Viability stain (1x PBS diluent, 20 min, room temperature) | | | | |
| LIVE/DEAD fixable blue viability dye (BViD) | UV450 | NA | Thermo Fisher | 1:500 |
| 1º Surface stain (FACSwash + Azide diluent, 20 min, room temperature) | | | | |
| CX3CR1 | PE-Cy7 | 2A9-1 | BioLegend | 1:160 |
| TruStain FcX (Fc block) | NA | NA | BioLegend | 1:20 |
| 2º Surface stain (FACSwash + Azide diluent, 20 min, room temperature) | | | | |
| Brilliant stain buffer concentrate | NA | NA | BD Biosciences | 1:5 |
| CD40 | BUV395 | 5C3 | BD Biosciences | 1:40 |
| CD16 | BUV496 | 3G8 | BD Biosciences | 1:320 |
| CD56 | BUV563 | NCAM16.2 | BD Biosciences | 1:160 |
| CD3 | BUV661 | UCHT1 | BD Biosciences | 1:80 |
| CD86 | BUV737 | 2331 | BD Biosciences | 1:40 |
| CD45 | BUV805 | HI30 | BD Biosciences | 1:80 |
| PD-L2 | BV421 | MIH18 | BD Biosciences | 1:20 |
| CD85k | BV480 | ZM3.8 | BD Biosciences | 1:40 |
| CD14 | BV570 | M5E2 | BioLegend | 1:20 |
| CD141 | BV605 | 1A4 | BD Biosciences | 1:640 |
| SIRPα | BV650 | SE5A5 | BD Biosciences | 1:160 |
| CD11b | BV750 | ICRF44 | BD Biosciences | 1:160 |
| CD123 | BV786 | 7G3 | BD Biosciences | 1:40 |
| CD206 | BB515 | 19.2 | BD Biosciences | 1:20 |
| BTLA | BB630 | J168-540 | BD Biosciences | 1:80 |
| PD-L1 | BB660 | MIH1 | BD Biosciences | 1:40 |
| CD32 | BB700 | FLI8.26 | BD Biosciences | 1:160 |
| CD38 | BB790 | HIT2 | BD Biosciences | 1:80 |
| MR1 | PE | 26.5 | BioLegend | 1:10 |
| CD137L (4-1BBL) | PE-Dz594 | 5F4 | BioLegend | 1:40 |
| CD80 | PE-Cy5 | L37.4 | BD Biosciences | 1:10 |
| CD19 | PE-Cy5.5 | SJ25-C1 | Thermo Fisher | 1:160 |
| CD1c | AF647 | F10/21A3 | BD Biosciences | 1:160 |
| CD11c | AF700 | B-ly6 | BD Biosciences | 1:320 |
| HLA-DR | APC-H7 | G46-6 | BD Biosciences | 1:40 |
| Fix (1x BD Cytofix/Cytoperm, 20 min, room temperature) | | | | |
| Intracellular stain (1x BD Perm/Wash buffer, 30 min, room temperature) | | | | |
| CD68 | BV711 | Y1/82A | BD Biosciences | 1:40 |

**Supplemental table 9**
**SA conjugates for HLA tetramerization and CyTOF screening**

| Reagent | Metal | Fluidigm metal kit |
|---|---|---|
| SA "1" | $^{155}$Gd | 201155B |
| SA "2" | $^{157}$Gd | Trace Sciences* |
| SA "3" | $^{159}$Tb | 201159B |
| SA "4" | $^{161}$Dy | 201161B |
| SA "5" | $^{163}$Dy | 201163B |
| SA "6" | $^{164}$Dy | 201164B |
| SA "7" | $^{167}$Er | 201167B |
| SA "8" | $^{170}$Er | 201170B |
| SA "9" | $^{173}$Yb | 201173B |
| SA "10" | $^{174}$Yb | 201174B |

*Isotope procured from Trace Sciences International

**Supplemental table 10**
**SA tetramer combinations for HLA-A*01, -A*02, -A*03, and/or -B*07 donors**

| SA combo. | HLA | Antigen | Peptide |
|---|---|---|---|
| $^{155}$Gd, $^{157}$Gd | A*01 | CMV UL44 | VTEHDTLLY |
| $^{155}$Gd, $^{159}$Tb | A*01 | CMV pp65 | YSEHPTFTSQY |
| $^{155}$Gd, $^{161}$Dy | A*01 | HCV polyprotein | ATDALMTGY |
| $^{155}$Gd, $^{163}$Dy | A*01 | IAV NP | CTELKLSDY |
| $^{155}$Gd, $^{164}$Dy | A*01 | IAV PB1 | VSDGGPNLY |
| $^{155}$Gd, $^{167}$Er | A*01 | AdV Hexon | TDLGQNLLY |
| $^{155}$Gd, $^{170}$Yb | A*01 | HSV1 UL46 (354–362) | ATDSLNNEY |
| $^{155}$Gd, $^{173}$Yb | A*01 | HSV1 UL48 (90–99) | SALPTNADLY |
| $^{155}$Gd, $^{174}$Yb | A*01 | HSV1 UL48 (479–488) | FTDALGIDEY |
| $^{157}$Gd, $^{159}$Tb | A*02 | CMV IE1 | VLEETSVML |
| $^{157}$Gd, $^{161}$Dy | A*02 | CMV pp65 | NLVPMVATV |
| $^{157}$Gd, $^{163}$Dy | A*02 | EBV LMP2A | CLGGLLTMV |
| $^{157}$Gd, $^{164}$Dy | A*02 | EBV BRLF1 | YVLDHLIVV |
| $^{157}$Gd, $^{167}$Er | A*02 | EBV BLMF1 | GLCTLVAML |
| $^{157}$Gd, $^{170}$Yb | A*02 | IAV M1 | GILGFVFTL |
| $^{157}$Gd, $^{173}$Yb | A*02 | EBV LMP1-2 | YLQQNWWTL |
| $^{157}$Gd, $^{174}$Yb | A*02 | EBV LMP1-1 | YLLEMLWRL |
| $^{159}$Tb, $^{161}$Dy | A*02 | CMV pp65-2 | QMWQARLTV |
| $^{159}$Tb, $^{163}$Dy | A*02 | Proinsulin precursor (15–24) | ALWGPDPAAA |
| $^{159}$Tb, $^{164}$Dy | A*02 | VZV IE62 (593–601) | ALWALPHAA |
| $^{159}$Tb, $^{167}$Er | A*02 | EBV BALF-4 | FLDKGTYTL |
| $^{159}$Tb, $^{170}$Yb | A*02 | EBV BMRF-1 | TLDYKPLSV |
| $^{159}$Tb, $^{173}$Yb | A*02 | MelanA/MART (26–35) | ELAGIGILTV |
| $^{159}$Tb, $^{174}$Yb | A*02 | MAGEA3 (271–279) | FLWGPRALV |
| $^{161}$Dy, $^{163}$Dy | A*02 | HSV1/2 UL25 (367–375) | FLWEDQTLL |
| $^{161}$Dy, $^{164}$Dy | A*02 | HSV1 UL47 (286-294) | FLADAVVRL |
| $^{161}$Dy, $^{167}$Er | A*02 | HSV1 UL47 (545–553) | RLLGFADTV |
| $^{161}$Dy, $^{170}$Yb | A*03 | EBV EMNA 3A | RLRAEAQVK |
| $^{161}$Dy, $^{173}$Yb | A*03 | HPV E6 | IVYRDGNPY |
| $^{161}$Dy, $^{174}$Yb | A*03 | CMV IE-1 | KLGGALQAK |
| $^{163}$Dy, $^{164}$Dy | A*03 | IAV PR8 | ILRGSVAHK |
| $^{163}$Dy, $^{167}$Er | A*03 | HSV RS1 (1096–1105) | RLYPDAPPLR |
| $^{163}$Dy, $^{170}$Yb | B*07 | CMV pp65 | RPHERNGFTVL |
| $^{163}$Dy, $^{173}$Yb | B*07 | CMV pp65 | TPRVTGGGAM |
| $^{163}$Dy, $^{174}$Yb | B*07 | EBV BMRF1 | RPQGGSRPEFVKL |
| $^{164}$Dy, $^{167}$Er | B*07 | EBV EBNA 3A | RPPIFIRRL |
| $^{164}$Dy, $^{170}$Yb | B*07 | EBV EBNA 6 | QPRAPIRPI |
| $^{164}$Dy, $^{173}$Yb | B*07 | IAV NP | SPIVPSFDM |
| $^{164}$Dy, $^{174}$Yb | B*07 | IAV PB1 | QPEWFRNVL |
| $^{167}$Er, $^{170}$Yb | B*07 | HSV ICP0 (698–706) | VPGWSRRTL |
| $^{167}$Er, $^{173}$Yb | B*07 | HSV1 UL21 (382–390) | VPRPDDPVL |
| $^{167}$Er, $^{174}$Yb | B*07 | HAV1 UL49 (281–290) | RPTERPRAPA |
| $^{170}$Yb, $^{173}$Yb | A*02 | SMCY/JARID1D | FIDSYICQV |
| $^{170}$Yb, $^{174}$Yb | B*07 | SMCY/JARID1D | SPSVDKARAEL |
| $^{173}$Yb, $^{174}$Yb | A*02 | CMV pp65 | NLVPTVAML |

**Supplemental table 11**
**SA tetramer combinations for HLA-A*02 and/or -A*11 donors**

| SA combo. | HLA | Antigen | Peptide |
|---|---|---|---|
| $^{155}$Gd, $^{157}$Gd | A*02 | PCDH11Y | FLLENAAYLD |
| $^{155}$Gd, $^{159}$Tb | A*02 | USP9Y | LLMPAGVPLT |
| $^{155}$Gd, $^{161}$Dy | A*02 | DDX3Y | FLLDILGAT |
| $^{155}$Gd, $^{163}$Dy | A*02 | UTY | LLIADNPQL |
| $^{155}$Gd, $^{164}$Dy | A*02 | UTY | YLQQNTHTL |
| $^{155}$Gd, $^{167}$Er | A*02 | UTY | ALIDCNPCTL |
| $^{155}$Gd, $^{170}$Yb | A*02 | UTY | KLLSSGAFSA |
| $^{155}$Gd, $^{173}$Yb | A*02 | UTY | LLSSGAFSA |
| $^{155}$Gd, $^{174}$Yb | A*02 | SMCY/JARID1D | FIDSYICQV |
| $^{157}$Gd, $^{159}$Tb | A*02 | SMCY/JARID1D | KMAAFPETL |
| $^{157}$Gd, $^{161}$Dy | A*02 | SMCY/JARID1D | SLMASSPTSI |
| $^{157}$Gd, $^{163}$Dy | A*02 | SMCY/JARID1D | HLLTSPKPSL |
| $^{157}$Gd, $^{164}$Dy | A*02 | SMCY/JARID1D | NVMPVLDQSV |
| $^{157}$Gd, $^{167}$Er | A*02 | SMCY/JARID1D | LMASSPTSI |
| $^{157}$Gd, $^{170}$Yb | NA | NA | NA |
| $^{157}$Gd, $^{173}$Yb | NA | NA | NA |
| $^{157}$Gd, $^{174}$Yb | NA | NA | NA |
| $^{159}$Tb, $^{161}$Dy | A*02 | CMV IE1 | VLEETSVML |
| $^{159}$Tb, $^{163}$Dy | A*02 | CMV pp65 | NLVPMVATV |
| $^{159}$Tb, $^{164}$Dy | A*02 | EBV LMP2A | CLGGLLTMV |
| $^{159}$Tb, $^{167}$Er | A*02 | EBV BRLF1 | YVLDHLIVV |
| $^{159}$Tb, $^{170}$Yb | A*02 | EBV BLMF1 | GLCTLVAML |
| $^{159}$Tb, $^{173}$Yb | A*02 | IAV M1 | GILGFVFTL |
| $^{159}$Tb, $^{174}$Yb | A*02 | EBV LMP1-2 | YLQQNWWTL |
| $^{161}$Dy, $^{163}$Dy | A*02 | EBV LMP1-1 | YLLEMLWRL |
| $^{161}$Dy, $^{164}$Dy | A*02 | CMV pp65-2 | QMWQARLTV |
| $^{161}$Dy, $^{167}$Er | A*02 | Proinsulin precursor (15–24) | ALWGPDPAAA |
| $^{161}$Dy, $^{170}$Yb | A*02 | VZV IE62 (593–601) | ALWALPHAA |
| $^{161}$Dy, $^{173}$Yb | A*02 | EBV BALF-4 | FLDKGTYTL |
| $^{161}$Dy, $^{174}$Yb | A*02 | EBV BMRF-1 | TLDYKPLSV |
| $^{163}$Dy, $^{164}$Dy | A*02 | MelanA/MART (26–35) | ELAGIGILTV |
| $^{163}$Dy, $^{167}$Er | A*02 | MAGEA3 (271–279) | FLWGPRALV |
| $^{163}$Dy, $^{170}$Yb | A*02 | HSV1/2 UL25 (367–375) | FLWEDQTLL |
| $^{163}$Dy, $^{173}$Yb | A*02 | HSV1 UL47 (286–294) | FLADAVVRL |
| $^{163}$Dy, $^{174}$Yb | A*02 | HSV UL47 (545–553) | RLLGFADTV |
| $^{164}$Dy, $^{167}$Er | A*02 | RCC c-MET (654–662) | YVDPVITSI |
| $^{164}$Dy, $^{170}$Yb | A*02 | RCC MUC1 (950–958) | STAPPVHNV |
| $^{164}$Dy, $^{173}$Yb | A*02 | RCC (146–154) (A02) | ALCNTDSPL |
| $^{164}$Dy, $^{174}$Yb | A*02 | RCC Survivin (96–104) | LTLGEFLKL |
| $^{167}$Er, $^{170}$Yb | A*02 | RCC EphA2 (58) | IMNDMPIYM |
| $^{167}$Er, $^{173}$Yb | A*11 | EBV EBNA 3B | AVFDRKSDAK |
| $^{167}$Er, $^{174}$Yb | A*11 | EBV EBNA 4 | IVTDFSVIK |
| $^{170}$Yb, $^{173}$Yb | A*11 | EBV BRFL1 | ATIGTAMYK |
| $^{170}$Yb, $^{174}$Yb | A*11 | EBV LMP2 | SSCSSCPLSK |
| $^{173}$Yb, $^{174}$Yb | A*11 | IAV H1N2 PB2 | KVYKTYFEK |

**Supplemental table 12**
**CyTOF tetramer screen staining panel**

| Reagent | Conjugate | Clone | Vendor | Metal | Fluidigm metal kit | Reagent dilution |
|---|---|---|---|---|---|---|
| Tetramer and 1º Surface stain (1x PBS + 0.5% FBS, 0.02% sodium azide; 60 min, room temperature) | | | | | | |
| TCRγδ | PE | 5A6.E9 | ThermoFisher | NA | NA | 1:200 |
| CD8 | Pure | RPA-T8 | BioLegend | $^{111}Cd$ | 201111A | 1:100 |
| CXCR3 | Pure | G025H7 | BioLegend | $^{158}Gd$ | 201158B | 1:200 |
| PD-1 | Pure | eBioJ105 | ThermoFisher | $^{160}Gd$ | 201160B | 1:200 |
| CXCR5 | Pure | RF8B2 | BD Biosciences | $^{165}Ho$ | 201165B | 1:100 |
| CCR7 | Pure | 150503 | R&D Systems | $^{168}Er$ | 201168B | 1:100 |
| CCR5 | $^{171}Yb$ | NP-6G4 | Fluidigm | $^{171}Yb$ | NA | 1:100 |
| Viability stain (1x PBS diluent, 10 min, on ice) | | | | | | |
| Cisplatin | NA | NA | Fluidigm | $^{194/195}Pt$ | NA | 1:8000 |
| 2º Surface stain (1x PBS + 0.5% FBS, 0.02% sodium azide; 20 min, on ice) | | | | | | |
| CD4 | Pure | RPA-T4 | BioLegend | $^{110}Cd$ | 201110A | 1:100 |
| CD14 | Pure | M5E2 | BioLegend | $^{112}Cd$ | 201112A | 1:50 |
| CD19 | Pure | HIB19 | BioLegend | $^{113}Cd$ | 201113A | 1:50 |
| CD56 | Pure | NCAM16.2 | BD Biosciences | $^{114}Cd$ | 201114A | 1:50 |
| CD45 | $^{89}Y$ | HI30 | Fluidigm | $^{89}Y$ | NA | 1:100 |
| CD57 | Pure | HNK-1 | BioLegend | $^{115}In$ | Trace Sciences* | 1:200 |
| CLA | Pure | HECA-452 | BioLegend | $^{141}Pr$ | 201141B | 1:200 |
| HLA-DR | Pure | L243 | BioLegend | $^{142}Nd$ | 201142B | 1:100 |
| ITB7 | Pure | FIB504 | BioLegend | $^{143}Nd$ | 201143B | 1:200 |
| TIGIT | Pure | 741182 | R&D Systems | $^{144}Nd$ | 201144B | 1:50 |
| PE | Pure | PE001 | BioLegend | $^{146}Nd$ | 201146B | 1:200 |
| CD69 | Pure | FN50 | BioLegend | $^{147}Sm$ | 201147B | 1:100 |
| CD45RO | Pure | UCHL1 | BioLegend | $^{148}Nd$ | 201148B | 1:200 |
| CD161 | Pure | HP-3G10 | BioLegend | $^{149}Sm$ | 201149B | 1:100 |
| KLRG1 | Pure | 13F12F2 | ThermoFisher | $^{150}Nd$ | 201150B | 1:200 |
| CD27 | Pure | LG.7F9 | ThermoFisher | $^{151}Eu$ | 201151B | 1:200 |
| CD137 | Pure | 4B4-1 | BioLegend | $^{152}Sm$ | 201152B | 1:100 |
| CD103 | Pure | B-Ly7 | ThermoFisher | $^{153}Eu$ | 201153B | 1:200 |
| CD3 | Pure | UCHT1 | BioLegend | $^{154}Sm$ | 201154B | 1:200 |
| TIM-3 | Pure | F38-2E2 | BioLegend | $^{162}Dy$ | 201162B | 1:200 |
| CD71 | Pure | CY1G4 | BioLegend | $^{166}Er$ | 201166B | 1:200 |
| CD45RA | Pure | HI100 | BioLegend | $^{169}Tm$ | 201169B | 1:200 |
| CD39 | Pure | A1 | BioLegend | $^{172}Yb$ | 201172B | 1:100 |
| CD38 | Pure | HIT2 | BioLegend | $^{176}Yb$ | 201176B | 1:100 |
| CD16 | $^{209}Bi$ | 3G8 | Fluidigm | $^{209}Bi$ | NA | 1:100 |
| Fix (2% PFA, overnight, 4ºC) | | | | | | |
| Intracellular stain (1x eBioscience FOXP3 permeabilization buffer, 30 min, room temperature) | | | | | | |
| Granzyme K | Pure | GM6C3 | Santa Cruz | $^{145}Nd$ | 201145B | 1:100 |
| CTLA-4 | Pure | BNI3 | BD Biosciences | $^{156}Gd$ | 201156B | 1:200 |
| Perforin | Pure | B-D48 | Abcam | $^{175}Lu$ | 201175B | 1:200 |
| DNA intercalator stain (1x PBS, 10 minutes, on ice) | | | | | | |
| Cell ID Ir | NA | NA | Fluidigm | $^{191/193}Ir$ | NA | 1:4000 |

*Isotope procured from Trace Sciences International

**Supplemental table 13**
**Placental immune infiltrate immunofluorescence panel**

| Tissue: placenta fixed in 1:4 BD Cytofix:1xPBS overnight | | | | |
|---|---|---|---|---|
| **Reagent** | **Conjugate** | **Clone** | **Vendor** | **Dilution** |
| 1º stain (1x TBS + 5% normal human serum diluent) 60 min, room temperature | | | | |
| Mouse anti-human CD8 | AF647 | RPA-T8 | BD Biosciences | 1:100 |
| Mouse anti-human CD163 | PE-CF594 | GHI/61 | BD Biosciences | 1:100 |
| Rabbit anti-human CD45 | Pure | EP322Y | Abcam | 1:100 |
| 2º stain (1x TBS + 5% normal human serum diluent) 60 min, room temperature | | | | |
| Donkey anti-rabbit IgG | Dy488 | Poly4064 | BioLegend | 1:200 |
| Nuclear intercalator stain (1x PBS diluent) 5 min, room temperature | | | | |
| DAPI | NA | NA | ThermoFisher | 1ng/mL |

**Supplemental table 14**
**Trp metabolite LC-MS standards**

| Metabolite name | Vendor | Molecular weight | Mass weighed (mg) | Molarity (mM) |
|---|---|---|---|---|
| Pool 1, dissolved in 15mL $H_2O$ | | | | |
| Indole | Sigma | 117.15 | 8.92 | 5.07611325 |
| Anthranilic acid | Sigma | 137.14 | 10.25 | 4.9827427 |
| Indoleacetic acid | Sigma | 175.18 | 13.08 | 4.97773718 |
| 5-Hydroxyindoleacetic acid | Sigma | 191.18 | 14.32 | 4.99354884 |
| L-Kynurenine | Sigma | 208.21 | 15.5 | 4.96293806 |
| Cinnabarinic acid | Sigma | 300.22 | 1.99 | 0.44189816 |
| Pool 2, dissolved in 9.08mL $H_2O$ | | | | |
| Nicotinic acid | Sigma | 123.11 | 5.54 | 4.95599187 |
| 3-Hydroxyanthranilic acid | Sigma | 153.14 | 6.96 | 5.00535343 |
| Serotonin (HCl salt) | Sigma | 212.68 | 9.66 | 5.00224117 |
| Indolepyruvate | Sigma | 203.19 | 9.16 | 4.96486329 |
| 5-Hydroxytryptophan | Sigma | 220.22 | 9.99 | 4.9960052 |
| Pool 3, dissolved in 15mL $H_2O$ | | | | |
| Picolinic acid | Sigma | 123.11 | 9.3 | 5.03614654 |
| Tryptamine | Sigma | 160.22 | 12.15 | 5.05554862 |
| Kynurenic acid | Sigma | 189.17 | 14.26 | 5.02546211 |
| L-Tryptophan | Sigma | 204.23 | 15.33 | 5.00416197 |
| 3-Hydroxykynurenine | Sigma | 224.2133 | 16.6 | 4.93577619 |
| Pool 4, dissolved in 9.08mL $H_2O$ | | | | |
| Trigonelline (HCl salt) | Sigma | 173.6 | 7.86 | 4.98639842 |
| Quinolinic acid | Sigma | 167.12 | 7.6 | 5.00840357 |
| Indole-3-propionic acid | Sigma | 189.21 | 8.59 | 4.999922 |
| Xanthurenic acid | Fisher | 205.17 | 9.28 | 4.9813639 |
| Melatonin | Sigma | 232.28 | 10.56 | 5.00686927 |
